# The interaction of the Arabidopsis Xyloglucan Xylosyltransferases XXTs with the COPII member SAR1 via their di-Arginine motifs is critical for delivery to the Golgi

**DOI:** 10.64898/2025.12.30.697048

**Authors:** Ning Zhang, Jordan D. Julian, Olga A Zabotina

**Affiliations:** Roy J Carver Department of Biochemistry, Biophysics and Molecular Biology, Iowa State University, Ames, IA 50011, USA

**Keywords:** Xyloglucan biosynthesis, Xyloglucan Xylosyltransferases, SAR1 protein, cargo selection by COPII complex, protein trafficking

## Abstract

Golgi-localized Xyloglucan Xylosyltransferases (XXT2 and XXT5) participate in xyloglucan biosynthesis, and to do this, they require the proper localization. The COPII complex is responsible for delivering cargo proteins from the ER to the Golgi, which is facilitated by the complex’s member proteins Sec24 and Sar1. Additionally, the N-termini of glycosyltransferases (GTs) play a crucial role in their transportation and localization. In this study, we demonstrated for the first time that XXTs interact with Sar1 protein in the COPII complex but not with Sec24, which was previously reported to be the main recruiter of cargo proteins into COPII-coated vesicles. The mutation of the arginine to glutamine residues of di-arginine motifs in the N-termini of XXT2 and XXT5 caused protein mislocalization and significantly reduced the strength of the interaction with Sar1. These mutations caused 90% of XXTs to either remain in the ER or localize to non-Golgi small compartments. In turn, such mislocalization significantly suppressed the recovery of xyloglucan biosynthesis in *Arabidopsis thaliana* (*Arabidopsis*) mutant plants (*xxt1xxt2* and *xxt3xxt4xxt5*), failing to restore their root phenotypes to normal. Our results demonstrate the interaction between cargo proteins and Sar1 proteins, highlighting the critical role of di-arginine motifs in this interaction. These results provide new insights into the mechanism of ER-to-Golgi delivery of plant GTs, which significantly advances our understanding of polysaccharide biosynthesis in the Golgi and the enzymes responsible for it.

**Significance statement:** This study demonstrates that plant glycosyltransferases directly interact with the SAR1 protein of the COPII complex. The di-arginine motifs present in the N-termini of glycosyltransferases play a critical role in cargo selection and transport from the ER to the Golgi apparatus via COPII-coated vesicles, while interacting with SAR1.

## Introduction

Xyloglucans (XyGs) belong to the hemicelluloses as the main component of the plant primary cell wall. XyGs are highly branched polysaccharides with a 1,4-β-linked glucan backbone branched with side chains composed of diverse monosaccharides (Scheller and Ulvskov 2010). Xyloglucan Xylosyltransferases (XXTs) transfer the xylosyl residues to the glucan backbone as the first step of the XyG side chain initiation during XyG biosynthesis in the Golgi (Cavalier et al. 2008; Zabotina et al. 2009; Zabotina 2012; Julian and Zabotina 2022; Zhang et al. 2023). XXTs are a subset of glycosyltransferases (GTs), a large family of enzymes responsible for the glycosylation of different molecules within the Golgi and ER. XXTs are type II membrane proteins and share the basic structure with most GTs, including the N-terminal cytosolic tail, transmembrane domain (TMD), stem region, and C-terminal catalytic domain (Culbertson et al. 2018; Julian and Zabotina 2022). The N-terminal cytosolic tail and TMD were shown to be important in the transportation and localization of the Golgi-localized GTs (Giraudo and Maccioni 2003; Schoberer et al. 2009, 2014, 2019a; Franke et al. 2013; Wang et al. 2013b; Becker et al. 2018; Zhang and Zabotina 2022). The ER-Golgi transportation and proper localization of GTs, including those involved in XyG biosynthesis, are critical prerequisites for reproducible and efficient polysaccharide biosynthesis.

The N-terminal cytosolic tails of GTs have been shown to play a key role in their trafficking from the ER to the Golgi in eucaryotic cells (Giraudo and Maccioni 2003; Schoberer et al. 2009; Franke et al. 2013; Wang et al. 2013b; Becker et al. 2018; Zhang and Zabotina 2022), where specific amino acid motifs are critical for their Golgi localization. For example, arginine residues in the cytosolic tail of *Arabidopsis thaliana* (*Arabidopsis*) xylosyltransferase (AtXylT) and *Nicotiana tabacum* (*N. tabacum*) N-acetylglucosaminyltransferase I (GnTI) were required for their recruitment to the Golgi (Schoberer et al. 2009). In addition, it was demonstrated that the specific motifs in the cytosolic tails of UDP-GalNAc:polypeptide N-acetylgalactosaminyltransferases (GalNAc-T1 and GalNAc-T2) and N-acetylglucosamine-1-phosphotransferase (PT) α/β-subunits are essential for their Golgi localization (Giraudo and Maccioni 2003; Franke et al. 2013; Wang et al. 2013b; Becker et al. 2018; Zhang and Zabotina 2022). The di-arginine motifs were identified in the N-termini of the Golgi-localized β-1,3-galactosyltransferase (GalT2), β-1,4-N-acetyl-galactosaminyltransferase (GalNAcT), GM3 sialyltransferase (Sial-T2), and β1,4-Galactosyltransferase (β1,4GT) proteins and the mutation of arginine residue to alanine altered their localization (Giraudo and Maccioni 2003).

The coat protein complex II (COPII)-coated cargo carriers mediate the trafficking path of proteins, including GTs, between the ER and the Golgi. This process has been well-studied in yeast and mammalian cells; however, studies in plants are limited (Zhang and Zabotina,2022). In *Arabidopsis*, the five small secretion-associated Ras-related GTPase 1 (Sar1) proteins, seven inner coat protein Sec23 isoforms, three inner coat protein Sec24 isoforms, two outer coat protein Sec13 isoforms, and two outer coat protein Sec31 isoforms have been found as homologs of the yeast proteins and are believed to function in COPII-coated vesicles. In *Arabidopsis*, AtSar1a localizes in the ER export sites (ERES) (Hanton et al. 2008) and has a protein-protein interaction with AtSec23a (Zeng et al. 2015). AtSar1a was involved in the ER export of the transcription factor bZIP28 under ER stress conditions (Zeng et al. 2015), and the overexpression of the GTP-restricted mutant of AtSar1a inhibited its ER-export (Hanton et al. 2008). In addition, it was demonstrated that AtSar1b also localizes at Golgi-associated ERES (Hanton et al. 2008; Wang et al. 2016) and forms the protein complex with OsSec23c (Wang et al. 2016). The RNA interference (RNAi) constructs of OsSar1a, OsSar1b, and OsSar1c, expressed simultaneously, suppressed the ER-export of storage proteins in the rice endosperm (Tian et al. 2013). AtSec23a, AtSec23d, AtSar1b, and AtSar1c were shown to be important for pollen development (Aboulela et al. 2018; Liang et al. 2020). In the moss *Physcomitrium* (*Physcomitrella*) *patens*, the KO mutant of PpSec23d caused ER morphology defects and reduced ER-to-Golgi trafficking of this protein (Liang et al. 2020). Similarly, AtSec24a was required to maintain ER morphology (Faso et al. 2009; Nakano et al. 2009), and its KO mutant showed a significant impact on plant growth and development (Faso et al. 2009). All three *AtSec24* isoforms are highly expressed in all *Arabidopsis* tissues and show a similar expression pattern and subcellular localization (Tanaka et al. 2013).

In yeast cells, sorting signal peptides at the protein N-/C-termini directly or indirectly interact with the COPII components in the coat vesicles. Sar1 proteins, part of the COPII complex, directly interact with cargo proteins (Kim et al. 2022; Xu et al. 2023), including GTs (Giraudo and Maccioni 2003; Srivastava et al. 2012), and their interactions can affect the assembly of COPII coat vesicles. The cytosolic tails of certain cargo GTs directly interact with Sar1 proteins and Sec23p, another component of the COPII complex, facilitating their trafficking (Giraudo and Maccioni 2003). For example, the direct protein-protein interaction between yeast Sar1 proteins and the synthetic cytosolic tails containing RR motifs of GalNAcT and GalT2 were observed *in vitro*. The mutation of RR to AA impaired this *in vitro* interaction (Giraudo and Maccioni 2003). The cytosolic tails of GalNAcT and GalT2 also interacted with Sec23p, and active Sar1 strengthened their protein-protein interaction (Giraudo and Maccioni 2003). Additionally, a single arginine residue (R11) in GnTI was sufficient for the protein localization to the Golgi. The arginine residue (R2) and lysine residue (K5) in its cytosolic tail (MRGYKFCCDFR) affected its ER-to-Golgi transportation (Schoberer et al. 2009).

In this study, we focused on two *Arabidopsis* GTs, XXT2 and XXT5, to investigate the role of their N-terminal cytosolic tails in ER-to-Golgi transportation. To reveal critical residues controlling XXTs’ transportation and potential interaction with COPII complex components, we selected XXT2 and XXT5 because they are the most critical for the biosynthesis of XyGs among the five known XXTs. In addition, they have different lengths of their cytosolic N-termini, with 22 residues in XXT2 and 44 residues in XXT5. The initial experiments using various truncated proteins revealed the short sequences, including di-arginine motifs, critical for their localization in the Golgi. The subsequent experiments with mutant proteins demonstrated the importance of these motifs for their interaction with Sar1 proteins in the COPII complex. The importance of the correct delivery of XXTs to the Golgi in order to function in XyG biosynthesis was confirmed using side-directed mutagenesis and complementation experiments in *Arabidopsis* KO mutant plants.

## Results

### Subcellular localization of truncated XXTs transiently expressed in *Arabidopsis* protoplasts

To investigate the function of the N-terminal cytosolic tail in the XXTs subcellular localization and transportation from ER to Golgi, the multiple deletion mutants of XXT2 and XXT5 were generated. Proteins with N-terminal cytosolic tails truncated to different lengths were transiently co-expressed with Golgi or ER marker proteins in *Arabidopsis* protoplasts. Observations of the subcellular localization of the truncated XXTs revealed specific protein sequences that influence their Golgi localization. As expected, the XXT2 with the full length of the N-terminus was localized in the Golgi, overlapping with the Golgi marker (Figure 1). The truncated XXT2Δ12A and XXT2Δ15A were also localized in the Golgi, indicating that the first 15 amino acids in the N-terminus of XXT2 did not impact its localization (Figure 1 & Figure S1). The truncation of the predicted full-length N-terminus of XXT2 altered its localization. The XXT2ΔN mutant displayed mainly three types of localization: ER localization, non-Golgi-dot localization, and aggregation in an unidentified location in the protoplasts (Figure 1). In some protoplasts, XXT2ΔN exhibited a mixture of two types of mislocalization: ER localization and non-Golgi dot localization. These results revealed that the specific sequence motif “**RALRQLK**” is critical for the XXT2 localization to Golgi. It is known that transmembrane domains (TMD) are essential for membrane protein localization (Hu et al. 2011; Srivastava et al. 2012; Wang et al. 2013a; Schoberer et al. 2014, 2019b; Wu et al. 2016; Yu and Zhang 2019). As anticipated, the deletion of a whole TMD of XXT2 resulted in the aggregation of XXT2ΔNM and XXT2ΔM in the protoplasts (Figure 1 & Figure S1A & S2).

**Figure 1.**
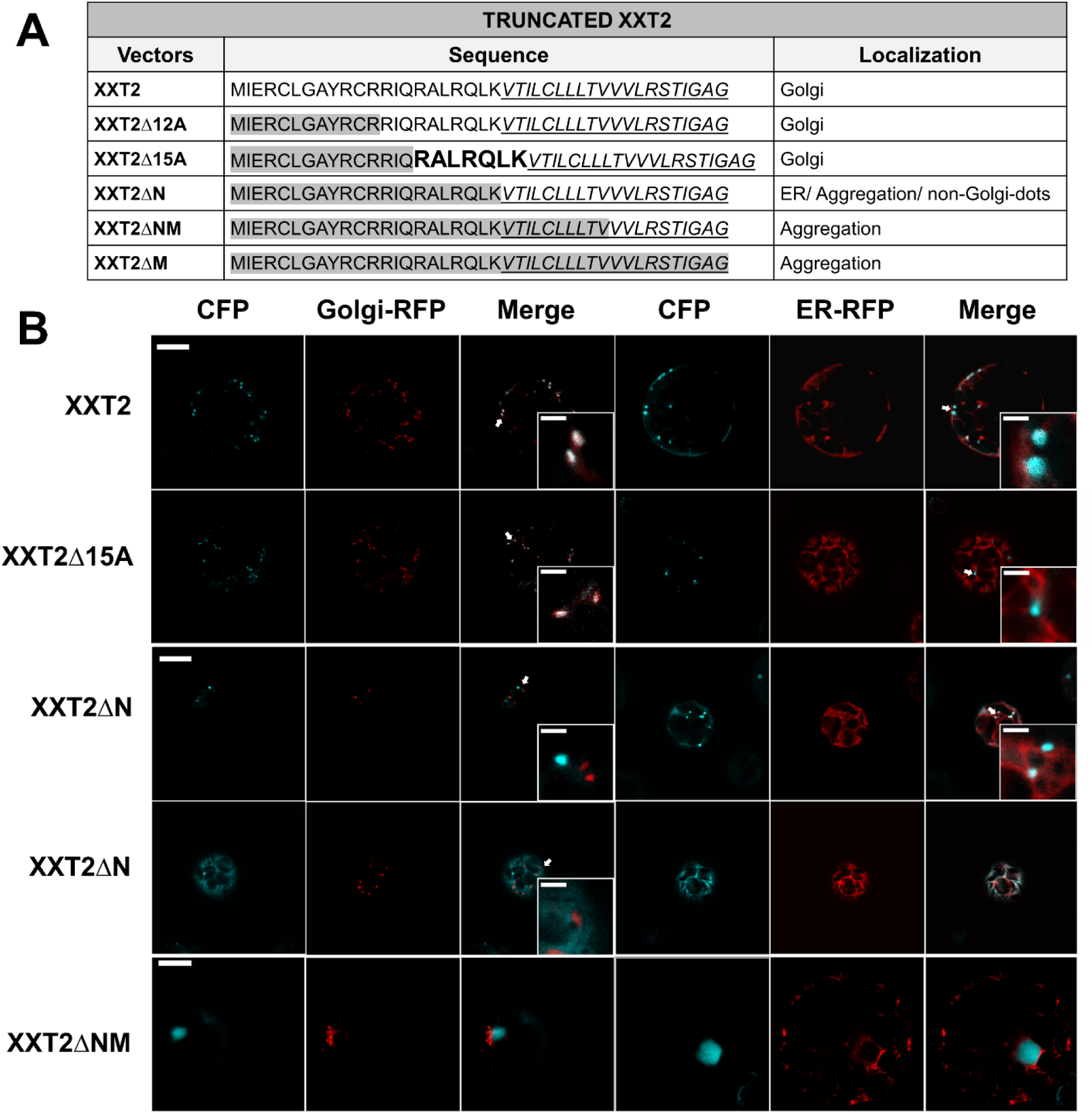
Subcellular localization of truncated XXT2 in Arabidopsis protoplasts. **(A)** The table summarizes the XXT2 deletion constructs used in this study; residues highlighted in gray represent truncated regions, italic underlined letters indicate predicted transmembrane domains, and the bold **RALRQLK** motif is required for Golgi localization of XXT2. **(B)** Subcellular localization of full-length XXT2 and truncation mutants co-expressed with Golgi-R (mCherry) or ER-R (mCherry) in Col-0 Arabidopsis protoplasts; The white arrows indicate Golgi-localized XXT2 and XXT2Δ15A and non-Golgi dot localization of XXT2ΔN. Scale bar = 10 μm; insets are shown at 4× magnification (scale bar = 2 μm).

Similar experiments using multiple deletion mutants were performed to investigate the importance of the XXT5 N-terminus for its localization. The truncation of the first 18 amino acids in the N-terminus of the XXT5 cytosolic tail did not show any impact on its Golgi localization; XXT5Δ10A and XXT5ΔTTT were localized in the Golgi (Figure 2 & Figure S1B & S3). The truncated XXT5 mutants (XXT5Δdi-Arg, XXT5Δ40A, and XXT5N), missing the specific sequence “**LPTTTLTNGGGRGGR**”, showed the three types of mis-localization: ER localization, the non-Golgi-dot localization, and aggregations (Figure 2 & Figure S1B & S3). These results demonstrated that the sequence “**LPTTTLTNGGGRGGR**” in the N-terminal cytosolic tail of XXT5 is critical for its Golgi localization.

**Figure 2.**
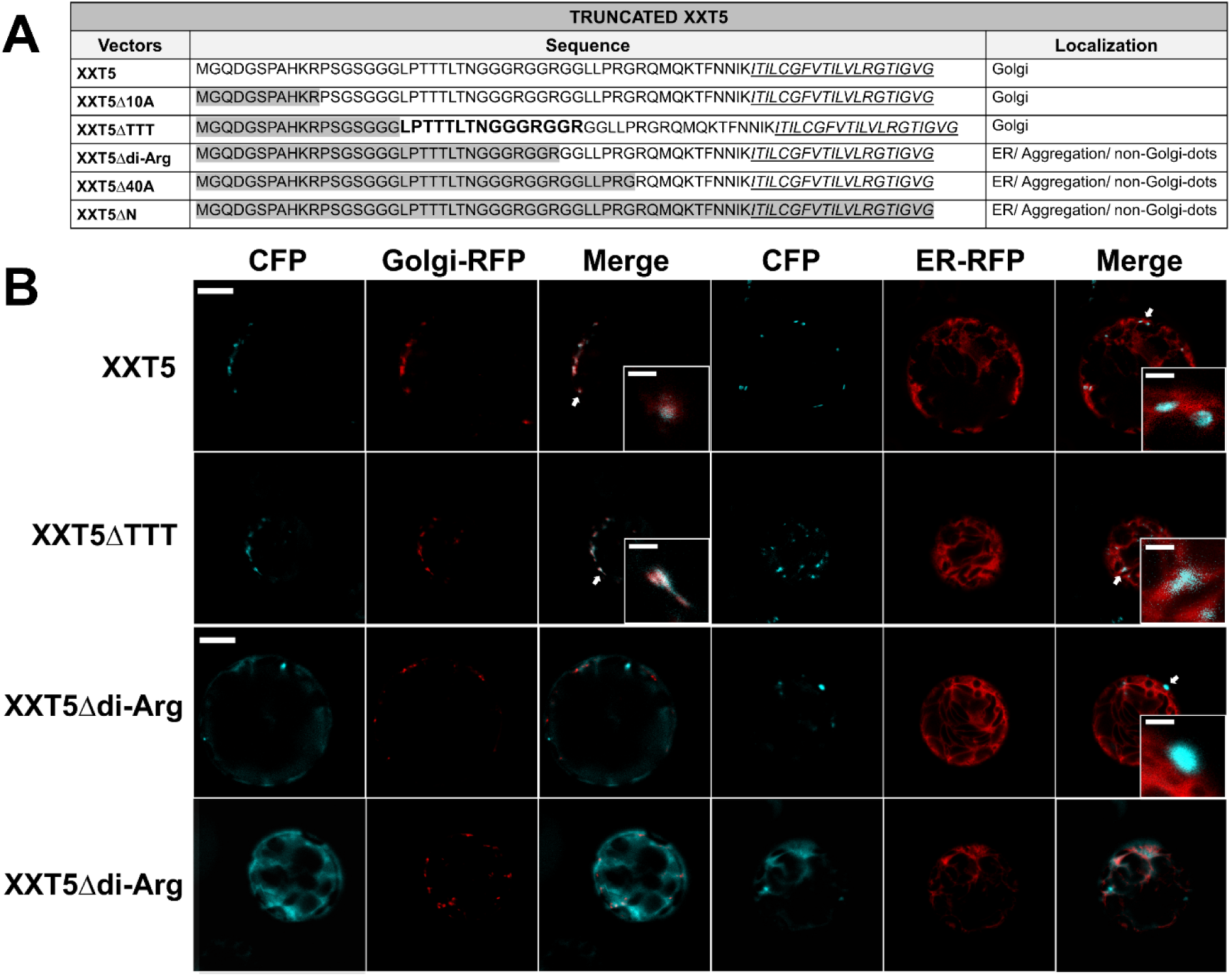
Subcellular localization of truncated XXT5 in Arabidopsis protoplasts. **(A)** The table summarizes the XXT5 deletion constructs used; residues highlighted in gray represent truncated regions, italic underlined letters indicate predicted transmembrane domains, and the bold **LPTTTLTNGGGRGGR** motif is required for Golgi localization of XXT5. **(B)** Subcellular localization of full-length XXT5 and mutants co-expressed with Golgi-R (mCherry) or ER-R (mCherry) in Col-0 Arabidopsis protoplasts; The white arrows indicate Golgi localization of XXT5 and XXT5ΔTTT and non-Golgi dot localization of XXT5Δdi-Arg. Scale bar = 10 μm; insets are shown at 4× magnification (scale bar = 2 μm).

### The di-arginine motif plays a critical role in determining the Golgi localization of XXT2 and XXT5

Earlier studies have reported that the di-arginine motifs play an essential role in the ER-Golgi transport of GTs and their localization(Giraudo and Maccioni 2003; Franke et al. 2013). The revealed sequences “**RALR**QLK” in XXT2 and “LPTTTLTNGGG**RGGR**” in XXT5 contain di-arginine motifs, as shown in bold. To confirm the function of these di-arginine motifs in the transportation of XXTs from ER to Golgi, the arginine residues were mutated to glutamine to maintain the sidechain structure but to eliminate the positive charge. The positive charges of amino acids are believed to be critical in protein-protein interactions(Lin and Chen 2004; Gymnopoulos et al. 2007) and can potentially impact trafficking processes. Various mutants in the N-terminus of XXT2 were generated. The mutant with the first arginine mutated to glutamine (**Q^16^**ALR^19^) was named XXT2-1RQ, the mutant with the second arginine mutated (R^16^AL**Q^19^**) was named XXT2-2RQ, and the mutant with both arginine residues mutated (**Q^16^**AL**Q^19^**) was named XXT2-RQRQ (Figure 3A). The transient expression of the XXT2-1RQ and XXT2-2RQ in the *Arabidopsis* protoplasts demonstrated that mutation of a single arginine in any position of the di-arginine motif resulted in mixed protein localization: both ER and Golgi (Figure 3 & Figure S4A & S5). The quantifications of XXT2-1RQ and XXT2-2RQ localization showed about 50% were ER-localized and another 50% were in Golgi or mixed localization in ER and Golgi in the same protoplast (Figure 3C). The mutation of both arginine residues resulted in mislocalization of the XXT2-RQRQ (Figure 3A), with 93.6% of the mutant protein localized either in the ER, non-Golgi-dots, or a mixture of two, whereas only 6.7% was found in the Golgi (Figure 3C & S4A).

**Figure 3.**
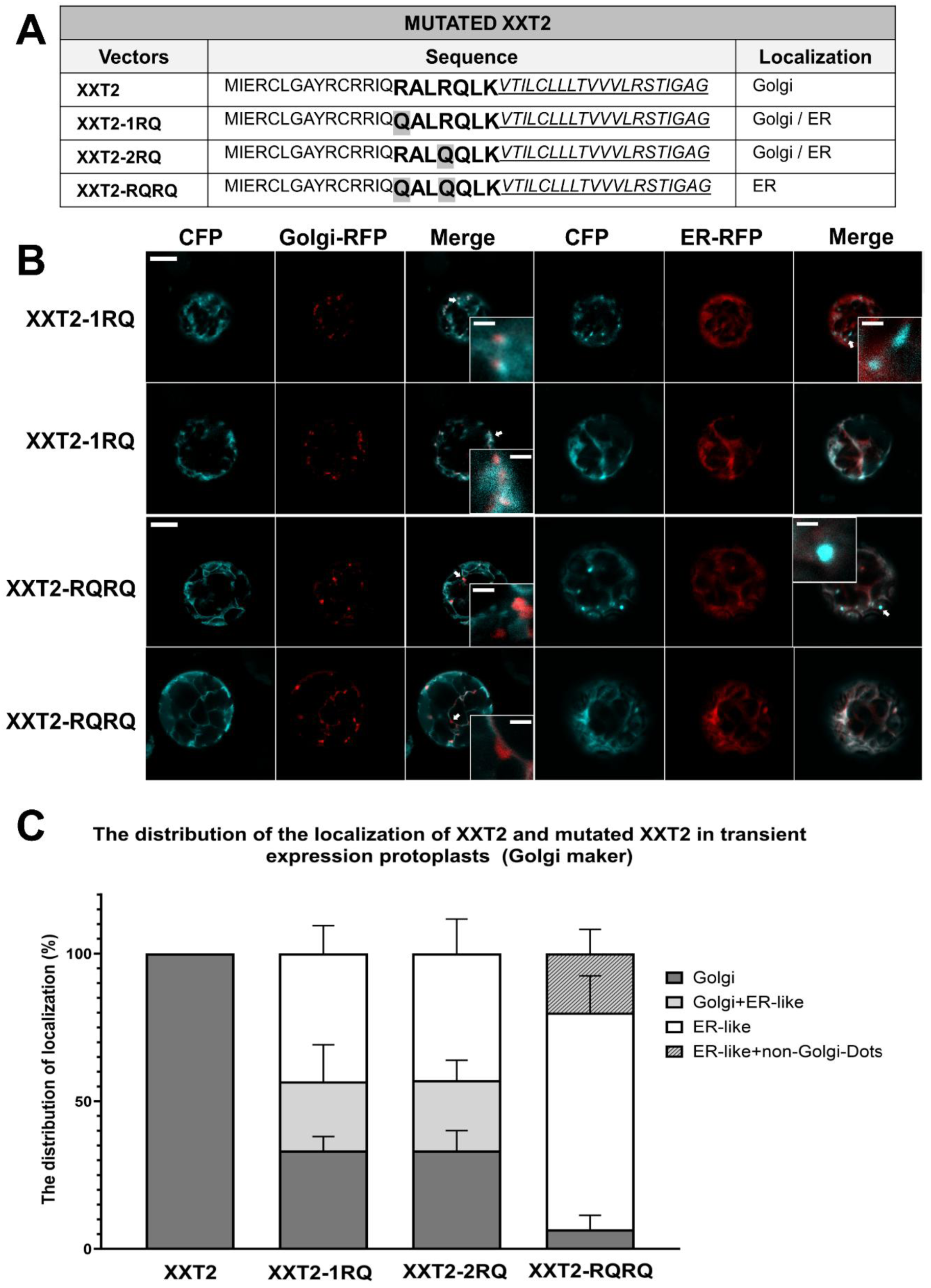
Subcellular localization of mutated XXT2 in Arabidopsis protoplasts. **(A)** The table summarizes the XXT2 mutant constructs; italic underlined residues indicate the predicted transmembrane region, and the bold **RALRQLK** motif is required for Golgi localization of XXT2. **(B)** Subcellular localization of XXT2 and mutants co-expressed with Golgi-R (mCherry) or ER-R (mCherry) in Col-0 protoplasts; scale bar = 10 μm, with insets shown at 4× magnification (scale bar = 2 μm). The white arrows indicate Golgi-dot localization of XXT2-1RQ and non-Golgi dot localization of XXT2-RQRQ. **(C)** Stacked bar graph showing quantitative analysis of the subcellular localization of XXT2 and mutant variants co-expressed with the Golgi marker in protoplasts; values represent mean ± SD from three independent experiments.

Similarly, one di-arginine motif is present in the sequence “LPTTTLTNGGG**R^30^GGR^33^**” of the cytosolic tail of XXT5, but another di-arginine motif (**R^39^GR^41^**) was found closer to its TMD (Figure 4A). To investigate the function of two di-arginine motifs in the XXT5 localization, the constructs of mutant proteins with both arginine residues mutated to glutamine in two separate di-arginine motifs were generated and named XXT5-RGGR and XXT5-RGR. The two arginine residues were both mutated to glutamine in the first di-arginine motifs **R^30^**GG**R^33^** in XXT5-RGGR, and two arginine residues were both mutated to glutamine in the second di-arginine motifs **R^39^**G**R^41^** in XXT5-RGR (Figure 4A). These mutant proteins, XXT5-RGGR and XXT5-RGR, were localized in both Golgi and ER (Figure 4 & Figure S4B & S6). The mutation of both di-arginine motifs (four arginine residues in total) in the XXT5-RQRQ (**Q^30^**GG**Q^33^**, **Q^39^**G**Q^41^**) mutant resulted in the three types of mislocalization of XXT5-RQRQ, similar to observed in the case of mutant XXT2: 87.18% of XXT5-RQRQ was localized either in the ER, non-Golgi-dots or a mixture of two, 10.27% of XXT5-RQRQ was found in Golgi and ER-like localizations, whereas only 2.56% was found in the Golgi (Figure 4C & S4B).

**Figure 4.**
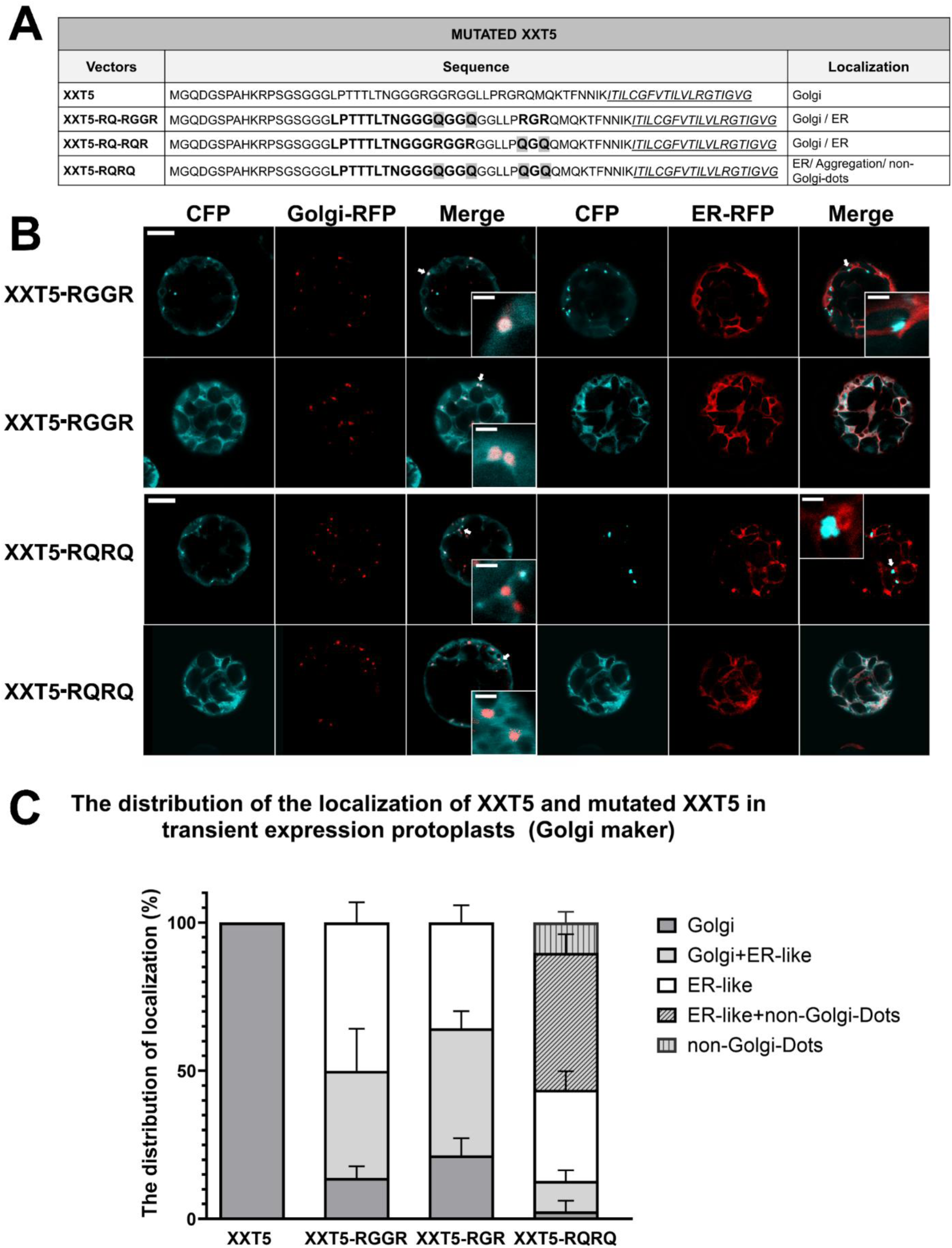
Subcellular localization of mutated XXT5 in Arabidopsis protoplasts. **(A)** The table summarizes the XXT5 mutant constructs; italic underlined residues denote the predicted transmembrane region, and the bold **LPTTTLTNGGGRGGR** motif is required for Golgi localization of XXT5. **(B)** Subcellular localization of XXT5 and its mutants co-expressed with Golgi-R (mCherry) or ER-R (mCherry) in Col-0 Arabidopsis protoplasts; scale bar = 10 μm, with insets shown at 4× magnification (scale bar = 2 μm). The white arrows indicate Golgi-dot localization of XXT5 RGGR and non-Golgi-dot localization of XXT5-RQRQ. **(C)** Stacked bar graph showing quantitative analysis of the subcellular localization of XXT5 and mutant variants co-expressed with the Golgi marker in protoplasts; data represent mean ± SD from three independent experiments.

### The root hair phenotype of the transgenic plants expressing mutated XXT2 and XXT5

The delivery of XXTs to Golgi is required for their involvement in XyG biosynthesis, and their mislocalization or retaining in the ER should have a significant impact on this process. To confirm this premise, the wild-type XXT2 and mutant XXT2-RQRQ were stably expressed in the *xxt1xxt2* double KO mutant plants (Cavalier et al. 2008), while wild-type XXT5 and mutant XXT5-RQRQ were expressed in the *xxt3xxt4xxt5* triple KO mutant plants (Zhang et al. 2023). Both KO double and triple mutants have somewhat similar short-root hair phenotypes. The expression of XXT2 in the *xxt1xxt2* double mutant restored the short root hairs to a length comparable with Col-0 (Figure 5A & C). The expression of XXT2-RQRQ only partially recovered the root hair length and the bubble-like extrusions at the root hairs’ tips in comparison with double mutant plants (Figure 5A & C). Similarly, the expression of wild-type XXT5 rescued the short root hair of the *xxt3xxt4xxt5* triple mutant plants to the level of Col-0 (Figure 5B and D), while the expression of XXT5-RQRQ only partially recovered the root hair length. (Figure 5B and D).

**Figure 5.**
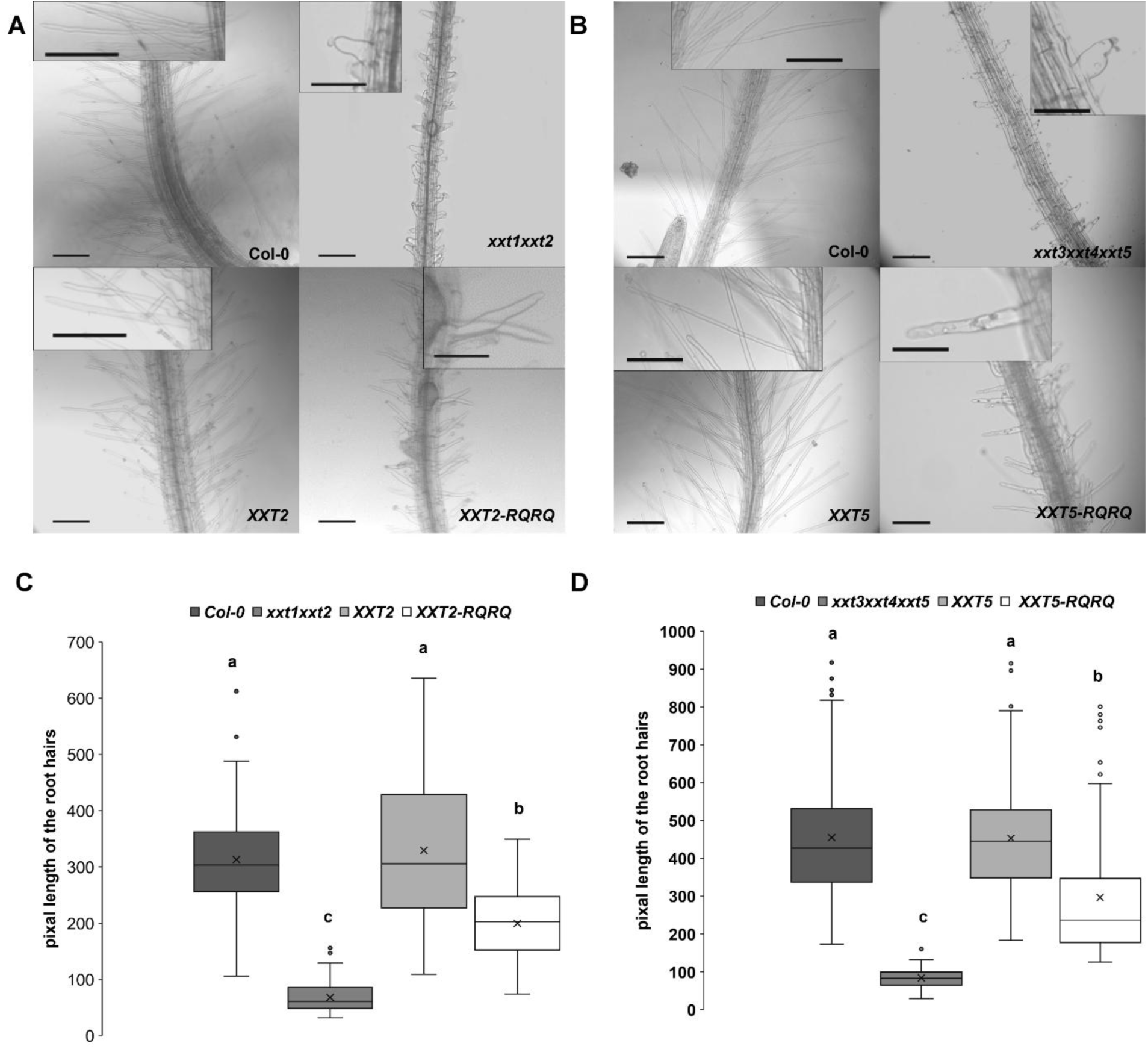
The Morphology and length of root hairs of transgenic Arabidopsis seedlings expressing XXTs and mutated XXTs. **(A)** Morphology and length of root hairs of Arabidopsis seedlings. Root hairs of 10-day-old Arabidopsis plants were observed under a microscope with 10 X magnification. The lengths of the root hairs of Col-0, XXT2, XXT2-RQRQ, and *xxt1xxt2*. Bar = 0.10 mm for all images. The insertion in Col-0 and XXT2 is enlarged two times (Bar = 0.10 mm). The insertion in XXT2-RQRQ and *xxt1xxt2* is enlarged three times (Bar = 0.05 mm). **(B)** The lengths of the root hairs of Col-0, XXT5, XXT5-RQRQ, and *xxt3xxt4xxt5*. Bar = 0.10 mm for all images. The insertion in Col-0 and XXT5 is enlarged two times (Bar = 0.10 mm). The insertion in XXT5-RQRQ and *xxt3xxt4xxt5* is enlarged three times (Bar = 0.05 mm). **(C & D)** Measurements in pixels from 15 independent seedlings for each line and more than 150 root hairs per line were used for statistical analyses. Different letters indicate significantly different values (one-way ANOVA, Tukey HSD, P <0.01).

### Subcellular localization of the mutant XXT proteins in the transgenic plants

The abnormal shape of root hairs of *Arabidopsis* KO mutants was only partially rescued by expressing XXT2-RQRQ in *xxt1xxt2* and the XXT5-RQRQ in *xxt3xxt4xxt5* transgenic lines. To investigate the localization of the mutant XXTs in the transgenic plants, protoplasts were isolated from these plants and used for imaging after transfection with Golgi and ER markers (Figure 6 & Figure 7 & Figure S7-S9). The images collected revealed that while 100% of the expressed wild-type XXT2 and XXT5 proteins were localized in the Golgi (Figure 6B & Figure 7B), only 11.11% of XXT2-RQRQ showed localization in the Golgi alone (Figure S7A), and 12.96% showed Golgi and ER-like mixed localization (Figure 6B). The other 75.93% were mis-localized and distributed among the non-Golgi dots and ER-like locations (Figure 6B). Similarly, only 20.69% of XXT5-RQRQ was delivered to the Golgi, including 8.57% localized to the Golgi alone (Figure S7B) and 12.12% showed mixed Golgi and ER-like localization, whereas the remaining 79.31% were distributed in ER-like and non-Golgi-dots locations (Figure 7B).

**Figure 6.**
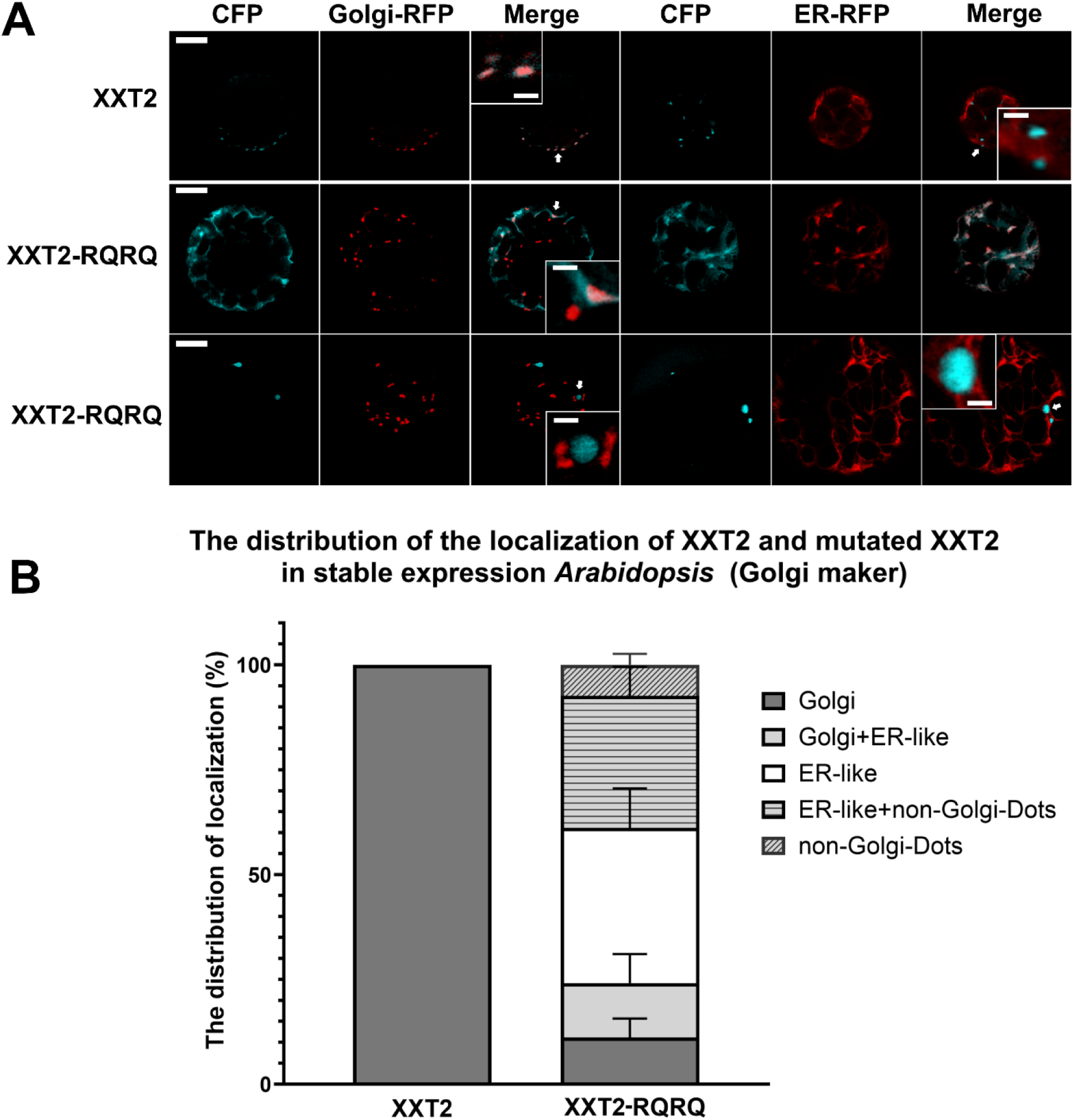
The subcellular localization of the wild type and mutated XXTs stably expressed in the Arabidopsis mutant plants. A: The subcellular localization of XXT2 and XXT2-RQRQ in the protoplasts prepared from the Arabidopsis *xxt1xxt2* plants expressing XXT2 or XXT2-RQRQ and transfected with Golgi-R (mCherry) marker. scale bar = 10 μm, with insets shown at 4× magnification (scale bar = 2 μm). The white arrows indicate the Golgi localization of XXT2; the non-Golgi dot localization of XXT2-RQRQ. B: The stacked bar figure shows the statistical analysis of the subcellular localization of XXT2 and XXT2-RQRQ with Golgi-maker in the protoplasts. Data means +/−SD from three replicate experiments.

**Figure 7.**
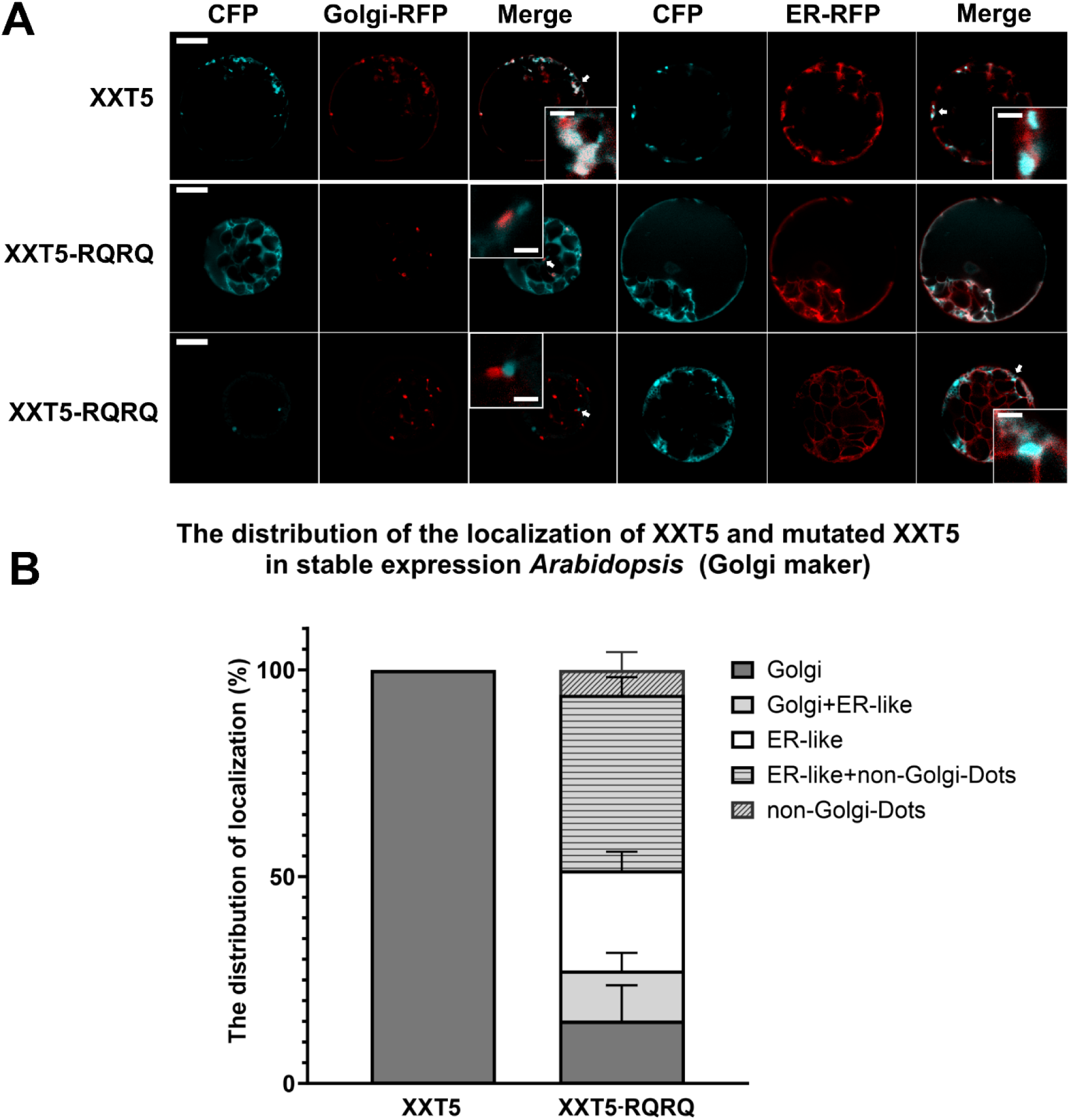
The subcellular localization of the wild type and mutated XXTs stably expressed in the Arabidopsis mutant plants. A: The subcellular localization of XXT5 and XXT5-RQRQ in the protoplasts prepared from the Arabidopsis *xxt3xxt4xxt5* plants expressing XXT5 or XXT5-RQRQ and transfected with Golgi R (mCherry) marker. scale bar = 10 μm, with insets shown at 4× magnification (scale bar = 2 μm). The white arrows indicate the Golgi localization of XXT5; the non-Golgi dot localization of XXT5-RQRQ. B: The stacked bar figure shows the statistical analysis of the subcellular localization of XXT5 and XXT5-RQRQ with Golgi-maker in the protoplasts. Data means +/−SD from three replicate experiments.

### The transgenic plants expressing mutant XXTs showed a significant reduction of XyGs in the cell walls

To investigate the structural patterns of the XyG molecules in all mutant and transgenic plants, the hemicellulose fractions were prepared from their cell walls, digested with the XyG-specific hydrolase, XEG, and analyzed by MALDI-TOF (Zabotina et al. 2012). As expected, XXXG, XXLG/XLXG, XXFG, and XLFG XyG subunits were observed in Col-0 (Figure 8D), whereas the *xxt1xxt2* mutant plant did not contain any detectable XyG (Cavalier et al. 2008) (Figure 8C). The *xxt3xxt4xxt5* triple mutant completely lacked the XLFG, XLLG, XXFG, and XXXG subunits, as was reported before (Zhang et al. 2023) (Figure 8G). All transgenic plants expressing either wild type or mutant XXT2 and XXT5 showed the subunit patterns of XyG somewhat similar to Col-0 plants, indicating that the synthesis of complete XyG was restored to a certain level (Figure 8 E, F and G).

**Figure 8.**
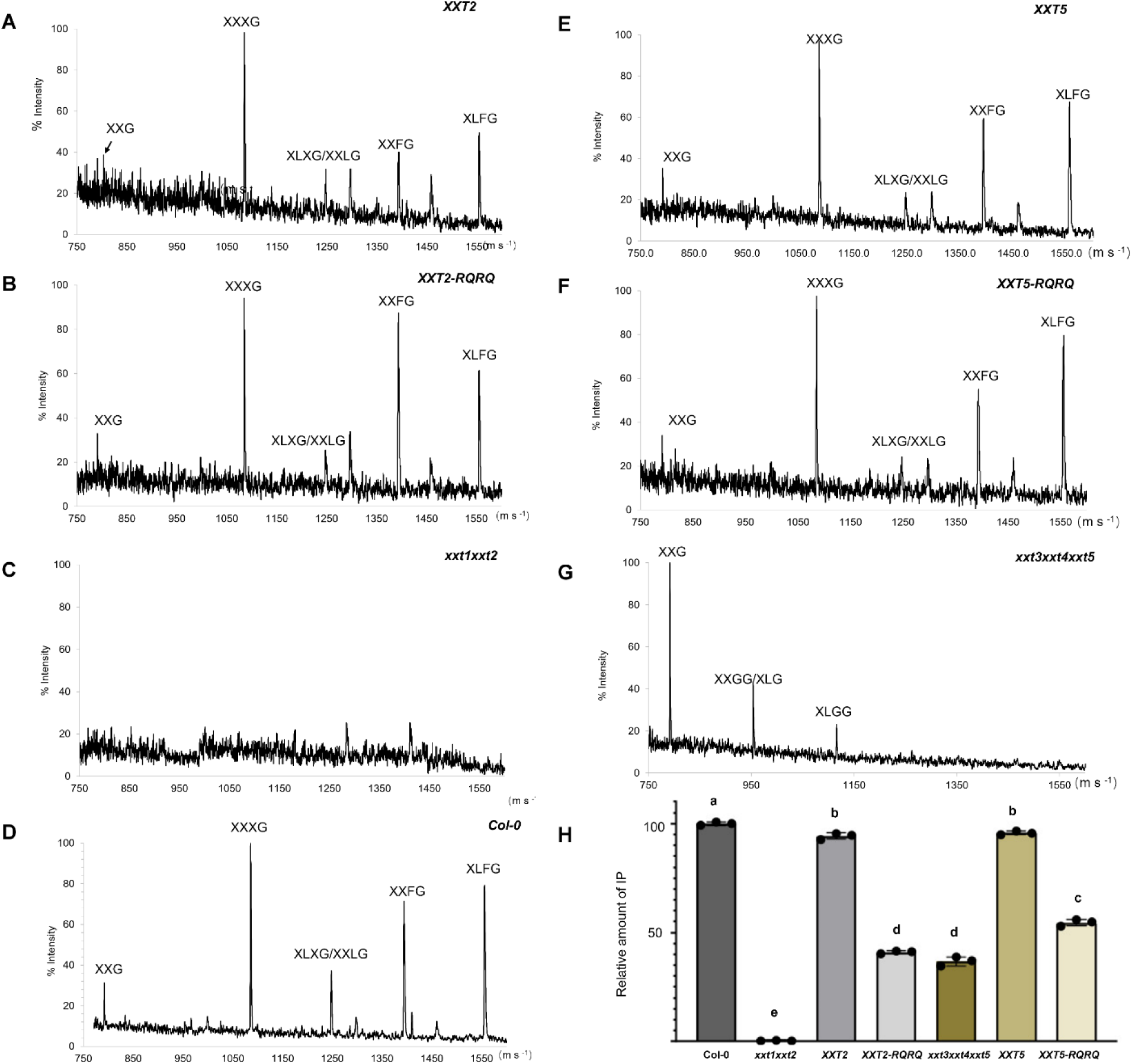
MALDI-TOF spectra of XEG-digested hemicellulose fractions extracted from all the mutants, transgenic lines, and Col-0 Arabidopsis plants. XXG-XLLG: xyloglucan subunits. A: XXT2 expressing in *xxt1xxt2* transgenic lines; B: XXT2-RQRQ expressing in *xxt1xxt2* transgenic lines; C: *xxt1xxt2*; D: Col-0; E: XXT5 expressing in xxt3xxt4xxt5 transgenic lines; F: XXT5-RQRQ expressing in *xxt3xxt4xxt5* transgenic lines; G: *xxt3xxt4xxt5*; H: Relative amounts of IP from the same hemicellulose fractions digested with Driselase and quantified using HPAEC analysis. The IP content obtained from the hemicellulose fraction from Col-0 was taken as 100. Data means +/− SD from three biological replicates, and different letters indicate significantly different values (one-way ANOVA, Tukey HSD, P <0.01).

Next, the total content of XyGs was determined in the cell walls prepared from all the above plants. Dry cell walls were digested with Driselase, and the amount of released signature disaccharide, isoprimeverose [IP; Xyl-a-(1-6)-Glc], was quantified by high-performance anion-exchange chromatography (HPAEC). The IP detected in all mutant and transgenic lines is expressed as a percentage relative to the amount of IP obtained from Col-0, which is set at 100% (Figure 8J). As expected, no IP was detected in the *xxt1xxt2* double mutants’ cell wall (Cavalier et al. 2008). In the *xxt3xxt4xxt5* mutant, the XyG content was decreased to approximately 40% (Figure 8J), as previously reported (Zhang et al. 2023). The plants expressing the wild type of XXT2 and XXT5 proteins showed almost complete restoration of XyG content to the level of Col-0 plants, while plants expressing mutant proteins had significantly decreased XyG content. Thus, the expression of XXT2-RQRQ in the *xxt1xxt2* plants restored the XyG content in their cell walls from 0% to 40% compared with the KO mutant and Col-0, respectively. The expression of XXT5-RQRQ in the *xxt3xxt4xxt5* plants increased XyG content by about 10% in comparison with the KO mutant (Figure 8J).

### The protein-protein interactions between XXTs and members of the COPII complex *in vivo* (BiFC) and *in vitro* pull-down assays

To understand whether XXTs are the cargo protein and interact with the main COPII-coated vesicle components, a BiFC assay was used to investigate the protein-protein interactions between XXTs and AtSar1/AtSec24a. BiFC signals were detected for nYFP-XXT2 with cYFP-Sar1d and for nYFP-XXT2 with cYFP-Sec24a (Figure 9 & Figure S10). Similarly, BiFC signals were also observed for cYFP-XXT5 with nYFP-Sar1d and for cYFP-XXT5 with nYFP-Sec24a (Figure 9). These results indicate that both AtSar1d and AtSec24a interact with XXT2 and XXT5. The mutant forms of XXT2 and XXT5 also retained their ability to interact with AtSar1d and AtSec24a (Figure 9).

**Figure 9.**
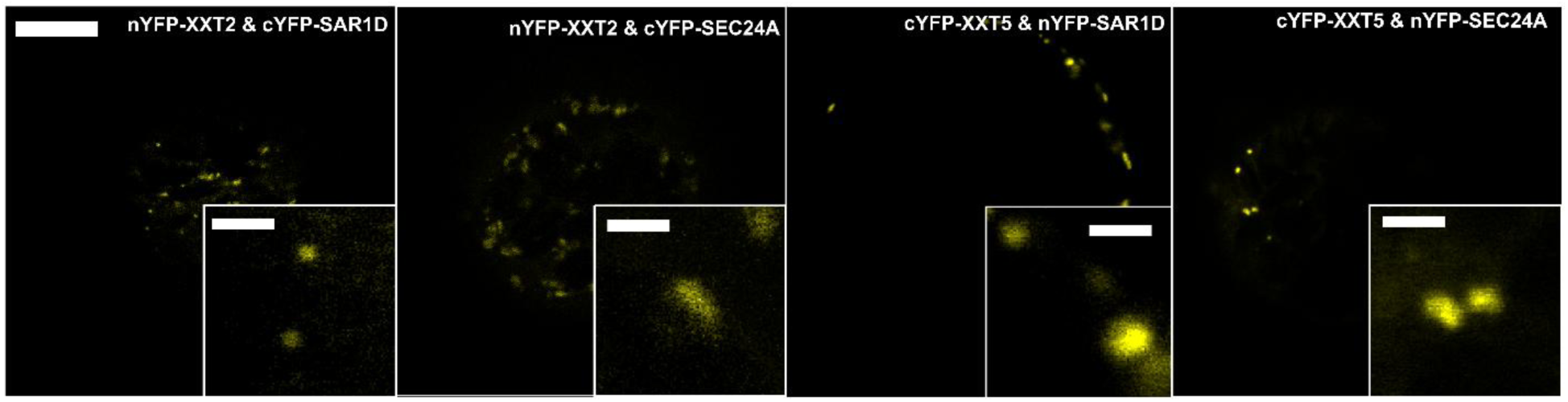
Fluorescence images of BiFC signal in Arabidopsis protoplasts. Bar, 10 μm with insets shown at 4× magnification (scale bar = 2 μm).

The Sar1 and Sec24a proteins are well known to form a heterodimer, and BiFC signals from their interaction with XXTs can arise simply from their dimeric arrangement. Therefore, we cannot distinguish whether XXTs interact with either Sar1 or Sec24a, or both, because the two COPII components are in close proximity. To separate these interactions, AtSar1 and AtSec24a were individually expressed in *E. coli* (BL21), and the recombinant proteins were extracted and purified for use as bait *in vitro* pull-down assays with full-length membrane proteins XXT2 and XXT5 obtained as total membrane protein extract from *Arabidopsis* plants. The primary anti-Myc antibody was used to detect AtSar1, the anti-HA antibody to detect AtSec24a, and the anti-GFP antibody to detect CFP-tagged XXT2 or XXT5. These experiments showed that full-length XXT2 and XXT5 interacted with Sar1b, Sar1c, and Sar1d, whereas no interaction with Sec24a was detected (Figure 10).

**Figure 10.**
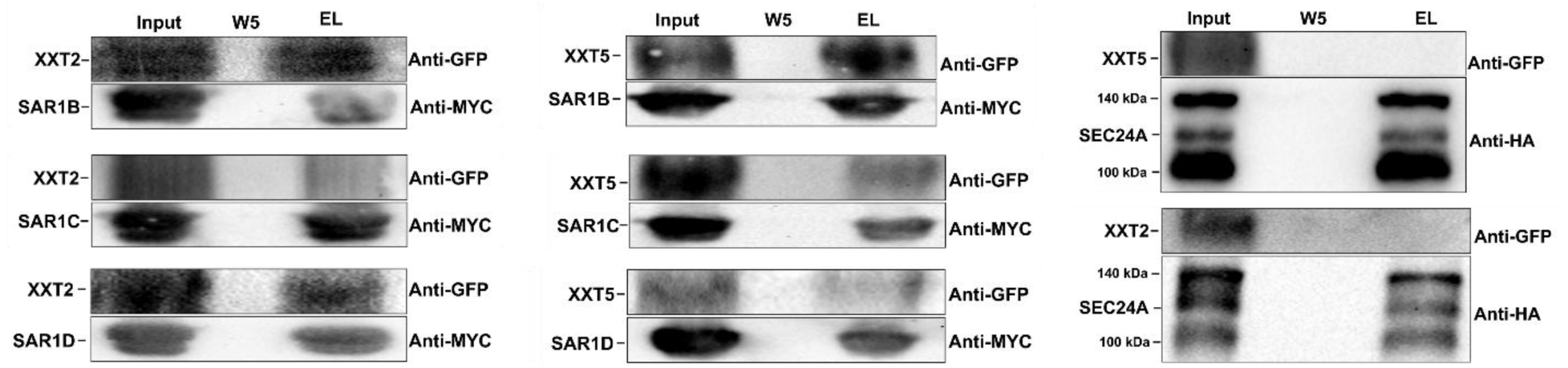
The *in vitro* pull-down assays demonstrating the protein-protein interactions between full length of XXTs with AtSar1 and AtSec24a proteins. A: Detection of the protein-protein interactions between XXT2 with AtSar1b, AtSar1c, and AtSar1d. B: Detection of the protein-protein interactions between XXT5 with AtSar1b, AtSar1c, and AtSar1d. C: Detection of the protein-protein interactions between XXT2 and XXT5 with AtSec24a protein.

XXTs are type II membrane proteins, with their cytosolic tails exposed to the cytosol and their stem regions and catalytic domains located in the Golgi lumen. This positioning makes the cytosolic tails the most likely region to interact with the soluble Sar1 protein. The peptides of N-terminal cytosolic tails of XXT2 and XXT5 were synthesized by the company (PEPTIDE 2.0). The wild-type peptides predicted to compose the cytosolic tails of wild-type XXT2 and XXT5 and the peptides with mutations R to Q (named XXT2Q) or from R to A (named XXT2A) were fused with a Flag tag. Two additional mutant peptides (R to A) were used to increase the potential impact of the mutation. First, it was confirmed that the amounts of wild-type peptides of XXTs and peptides with mutations used in the following pull-down experiments were comparable (Figure 10B-E, shown in the line corresponding to each peptide). Since the AtSar1 proteins lacked a Flag tag, they did not show any non-specific interaction with anti-Flag resin and served as the negative controls (Figure 10B-E). The COPII coat protein AtSar1b showed protein-protein interactions with the cytosolic tails of wild-type XXT2 and XXT5 (Figure 10B and C). The mutation from R to Q in the di-arginine motifs significantly affected the protein-protein interactions between the cytosolic tails of XXT5 and AtSar1b (Figure 10C) but did not affect the interaction between XXT2 and AtSar1b (Figure 10B). However, the peptides with R to A mutations showed significantly weaker interaction with AtSar1b for both XXT2 and XXT5 (Figure 10B and C). The peptides of both XXTs also interacted with AtSar1c and AtSar1d (Figure 10D-G). However, the strength of interactions was somewhat weaker than their interaction with Sar1b. Both types of mutations, either R to Q or R to A, significantly decreased the protein-protein interactions between both XXTs and AtSar1c or AtSar1d (Figure 10D-G). The experiments further confirmed the interactions between XXT peptides and AtSar1 protein, with no interactions observed with Sec24a (Figure S11).

## Discussion

Correct subcellular localization of proteins is the prerequisite for their normal function in the cells. Golgi-localized GTs must be delivered and retained in the Golgi after being properly folded in the ER. The importance of N-terminal cytosolic tails and TMDs of GTs for their transport and localization has been reported earlier (Giraudo and Maccioni 2003; Schoberer et al. 2009, 2014, 2019a; Franke et al. 2013; Wang et al. 2013b; Becker et al. 2018; Zhang and Zabotina 2022). In this study, we investigated the contribution of cytosolic tails and TMDs to the GTs localization in plant Golgi using two XXTs involved in plant polysaccharide biosynthesis. The deletion of N-terminal cytosolic tails of XXT2 and XXT5 altered their Golgi-localization (Figure 1, 2 & Figure S1), indicating the possible critical signaling role of their N-termini in ER-to-Golgi trafficking. Furthermore, deletion of both the N-terminus and TMD of XXT2 caused protein aggregation (Figure 1 & Figure S1), indicating the indispensable role of TMDs in proper GT trafficking via the secretory pathway.

By gradually deleting the cytosolic tails of XXTs, we identified the specific protein sequences containing di-arginine motifs as critical to their subcellular localization. Subsequent confirmation using the mutation of arginine residues showed that di-arginine motifs are primarily responsible for the proper Golgi localization of XXTs. Previous studies have reported the function of arginine residues and the di-arginine motifs in other proteins’ localization and trafficking (Giraudo and Maccioni 2003; Boulaflous et al. 2009; Schoberer et al. 2009; Uemura et al. 2009; Srivastava et al. 2012; Wang et al. 2013b; Zhang and Zabotina 2022). In our study, mutation from R to Q in the di-arginine motifs resulted in three types of mislocalization of 90% of mutated XXTs: ER-like, non-Golgi dots localization, and aggregation (Figure 3, 4 and Figure S4). This mislocalization of XXTs impeded the enzymes’ function in XyG biosynthesis. The XyG content in the transgenic plants expressing mutated XXTs was significantly reduced compared with Col-0 and KO mutants expressing corresponding wild-type XXTs (Figure 8). Moreover, expression of mutated XXTs resulted in significantly shorter root hairs compared to Col-0 plants (Figure 5). These results stipulate the critical role of the di-arginine motif in XXTs protein trafficking and function.

Notably, the double arginine residues in XXT2 and the double di-arginine motifs in XXT5 showed a cooperatively cumulative effect in the localization of proteins. The XXT2-1RQ and XXT2-2RQ with one random arginine residue mutation in the di-arginine motif triggered the mislocalization: about 76% of the mutated XXT2 localized in the ER and ER-Golgi mixed localization (Figure 3). The double mutation in both arginine residues caused higher mislocalization, with only 7% of XXT2-RQRQ remaining to be localized in the Golgi (Figure 3). Two di-arginine motifs were found in the cytosolic tail of XXT5, and both exerted influence on the subcellular localization of XXT5. The mutation in the four arginine residues of two di-arginine motifs significantly aggravated the mislocalization of XXT5-RQRQ (Figure 4). Similarly, our observation is consistent with previous findings that the mutation of any single arginine residue in the di-arginine motifs of Golgi-localized GalNAcT causes a mixed distribution (mainly in the Golgi and ER) (Giraudo and Maccioni 2003), whereas the mutation of both arginine residues results in ER localization.

In our study, we also observed the mutant XXTs in non-Golgi dot structures. This type of localization was identified due to the lack of overlap between the fluorescent signals of the Golgi marker and mutant XXTs; however, some of the XXT signals showed visible adherence to the ER marker and were surrounded by the ER (Figures 1–4). We speculate that the mislocalized mutant XXTs were retained in the ER, specifically at ER exit sites (ERESs). The COPII-coated vesicles are assembled in the ERESs (Rossanese et al. 1999; Hammond and Glick 2000; Ward et al. 2001; Brandizzi and Barlowe 2013). Cargo protein sorting, a critical step in protein trafficking, occurs during the assembly of COPII vesicles at ERESs. The plant homologous proteins Sec16 and MAG5 have been reported to localize at ERESs (Takagi et al. 2013, 2020). In that study, MAG5 was surrounded by a cup-shaped subdomain of the ER (Takagi et al. 2020). In our study, the non-Golgi dot localization of mutant XXT5-RQRQ showed a similar type of localization (Figure 4B). In the future, it would be interesting to co-express ERES marker MAG5 and mutant XXTs to understand the precise locations of mutant XXTs observed as non-Golgi dots.

Although mutation of the di-arginine motif altered the localization of the majority of XXTs, a small percentage of the XXT2-RQRQ and XXT5-RQRQ mutant proteins were still found in the Golgi; less than 10% of XXT2-RQRQ and 5% of XXT5-RQRQ were still delivered to the Golgi when transiently expressed in *Arabidopsis* protoplasts (Figure 3 and 4). A slightly higher content of mutant proteins was observed in the Golgi when constitutively expressed in KO plants; about 11% of XXT2-RQRQ and less than 10% of XXT5-RQRQ (Figures 6 and 7). The Golgi-localization of XXT is required for the successful transfer of the xylose residue on the glucan backbone during XyG biosynthesis. The MALDI-TOF results showed that matured, completely branched xyloglucan structures were synthesized in all transgenic plants, including XXTs with mutated arginine residues (Figure 8B, D, and F). This could indicate that as low as 10% of enzymes still delivered to the Golgi is sufficient to support some level of XyG synthesis. Such partial restoration of XyG in mutant cell walls was also sufficient for the partial restoration of root hair phenotypes normally observed in KO plants (Figure 5A and B). Since the substitution of both arginine residues does not entirely halt the XXT delivery to the Golgi, it is plausible that additional residues/motifs might contribute to this process. For example, it has been previously shown that the cooperation of N-termini with the TMDs/catalytic domain of GTs can contribute to the localization and transport of GTs (Franke et al., 2013; Wang et al., 2013b; Becker et al.,2018). Most likely, the mutation of the di-arginine motif in the cytosolic tails of XXTs significantly slowed down the rate of their delivery but did not eliminate their presence in the Golgi. For example, in yeast, it has been demonstrated that impairing the ER-export signal significantly decreases the transport rate and impedes the export of cargo proteins to the Golgi (Giraudo and Maccioni, 2003; Xu et al., 2023).

The COPII-coated vesicles carry the cargo from the ER to the Golgi in yeast, mammals, and plants. Specific motifs in cargo proteins have been reported to interact with components of the COPII-coat complex, including the Sec24 and Sar1 proteins. In yeast and mammalian cells, the Sec24 protein functions in cargo selection through direct protein-protein interactions with cargo proteins or cargo receptors (Miller et al., 2002, 2003; Mossessova et al., 2003; Wendeler et al., 2007; Pagant et al.,2015). Though the BiFC assay provided signals suggesting that XXTs interact with both Sar1 and Sec24A (Figure 9), this technique has limitations. Because Sar1 and Sec24a form a heterodimer, their close spatial proximity means that XXTs bound to either partner may bring the split fluorophore fragments together, producing fluorescence even when one of the apparent “interactions” is indirect. This spatial coupling can therefore generate false-positive signals that do not necessarily reflect direct binding between XXTs and either of the COPII components. Therefore, the result of BiFC should be corroborated by independent approaches, for example, pull-down experiments.

To avoid the impact of the Sar1–Sec24a heterodimer formation on the interpretation of results, AtSar1 and AtSec24a were expressed separately in *E. coli* BL21 as Myc- or HA-tagged recombinant proteins. The total membrane protein extracts from young *Arabidopsis* seedlings were used to pull down the full-length XXT2 and XXT5 proteins. These pull-down experiments showed that XXTs interact with Sar1 but not with Sec24a (Figure 10). These results strongly suggest that XXTs interact directly with Sar1 proteins, rather than with Sec24, as could be indicated by the BiFC assay. To further understand which region of XXTs mediates these interactions, it is important to consider that XXTs are type II membrane proteins, with their N-tails exposed to the cytosol and their stem regions and catalytic domains located in the Golgi lumen, making the cytosolic tails the most likely region to interact with soluble Sar1 and Sec24a protein. Synthetic peptides of XXT cytosolic tails were used in pull-down assays, which confirmed that these tails interact with Sar1 and not with Sec24a (Figure S11).

In *Arabidopsis*, there are five isoforms of Sar1 proteins, and it was proposed that their functions do not completely overlap (Hanton et al. 2008; Zeng et al. 2015, 2021; Feng et al. 2017; Liang et al. 2020; Li et al. 2021; Kim et al. 2022; Zhou et al. 2023). Here, we studied the role of AtSar1b, AtSar1c, and AtSar1d in the potential interaction with XXTs. These three Sar proteins were selected because the tissue-specific gene expression of AtSar1b, AtSar1c, and AtSar1d overlaps with that of XXTs (https://www.arabidopsis.org/), whereas AtSar1a is highly expressed only in siliques, and AtSar1e is highly expressed during seed germination (https://www.arabidopsis.org/). Results from BiFC, peptide-based pull-down assays, and full-length protein pull-down assays consistently demonstrated that XXTs interact with Sar1, including Sar1b, Sar1c, and Sar1d (Figures 9–11).

**Figure 11.**
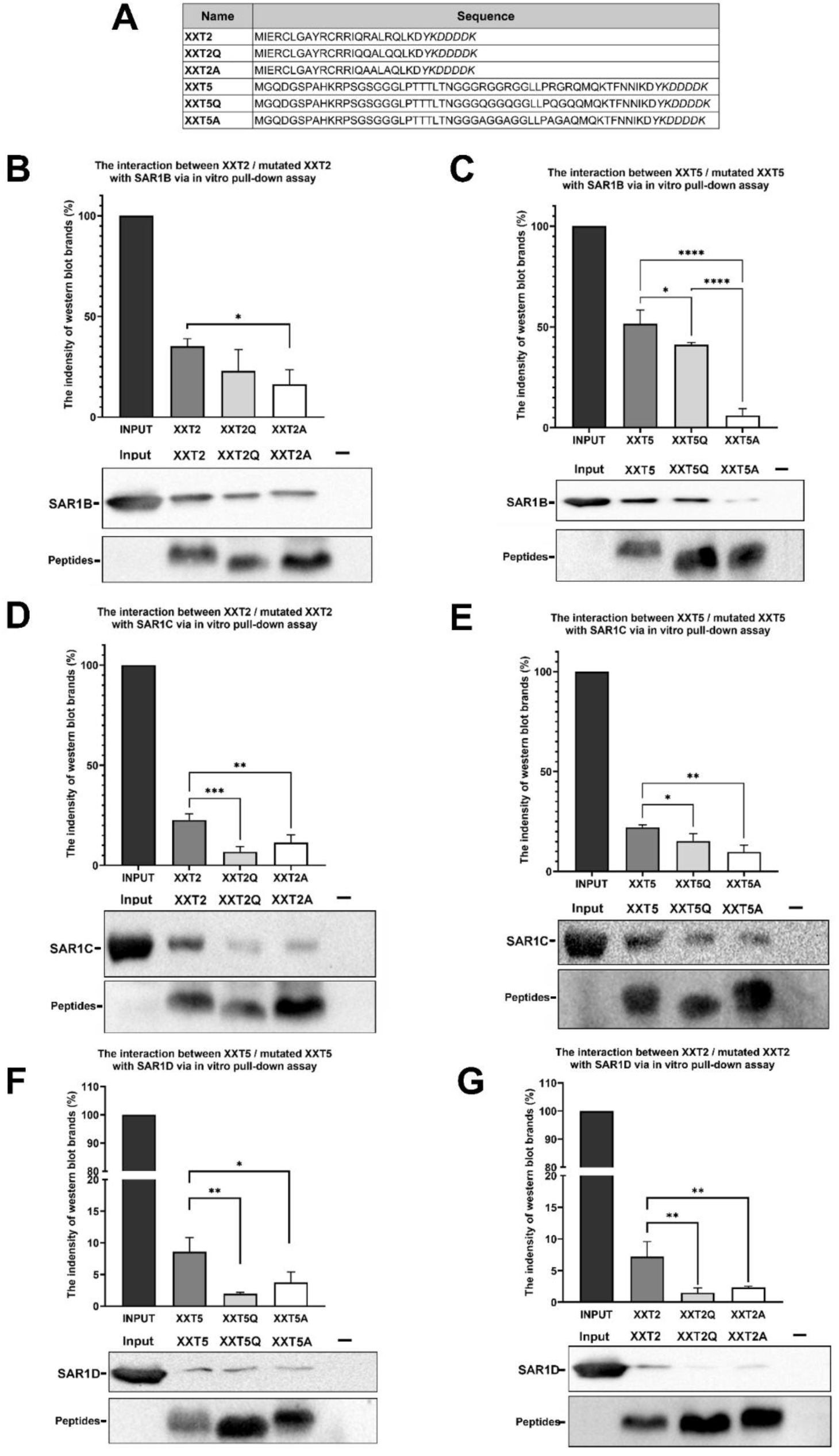
The in vitro pull-down assays demonstrate the protein-protein interactions between the cytosolic tails of XXTs and their mutants with AtSar1 proteins. A: The protein sequences of cytosolic tails of XXTs and their mutants. The Flag tag is labeled in italic font. Detection of the protein-protein interactions between XXT2 with AtSar1b (B), AtSar1c (D), and AtSar1d (F). Detection of the protein-protein interactions between XXT5 with AtSar1b (C), AtSar1c (E), and AtSar1d (G). The amount of peptides binding to the anti-Flag resin is shown in the peptide line, confirming that comparable amounts of peptides were bound to the anti-Flag resin. In the B, D, and F, the western blot results of XXT2, XXT2Q, and XXT2A peptides bound to an anti-Flag resin. In the C, E, and G, the western blot results of XXT5, XXT5Q, and XXT5A peptides bound to an anti-Flag resin. The band intensities were quantified using ImageJ (Fiji), and the Data Are Presented as means ± SD from three independent experiments. The asterisks indicate P<0.05, double asterisks indicate p<0.01, three asterisks indicate p<0.0005 and four asterisks indicate p<0.0001 in the unpaired t-test.

However, the XXTs exhibited different levels of interactions with AtSar1b, AtSar1c and AtSar1d, showing stronger interactions with AtSar1b compared to AtSar1c and AtSar1d (Figure 10). The substitution of R to Q residues in the di-arginine motif in cytosolic tails of XXTs weakened their interactions with AtSar1, and the substitution of R to A decreased interaction even further (Figure 10). The obtained difference between Sar1b, Sar1c, and Sar1d in their ability to interact with XXTs may imply that different isoforms of Sar1 might have diverse contributions to trafficking GTs in plants. The di-arginine motifs or [RK](X)[RK] motifs are found in the cytosolic tails of XXT1, XXT2, XXT5, XLT2, MUR3, and FUT1, the other GTs involved in XyG biosynthesis. Based on previous studies on the function of the SAR1 protein in cargo sorting (Giraudo and Maccioni 2003; Schoberer et al. 2009; Srivastava et al. 2012; Xu et al. 2023; Zhou et al. 2023) and our data from this study, we hypothesize that the Sar1 proteins are the primary players in the ER-to-Golgi trafficking of GTs involved in polysaccharide synthesis. Our results demonstrate that plant Golgi-resident membrane proteins such as GTs interact with the COPII complex through Sar1 proteins, and arginine residues in their N-terminal cytosolic tails are critical for this interaction, which determines the effectiveness of their delivery from the ER to the Golgi.

## Experimental procedures

### Plant growth and plasmid construction

The *xxt1xxt2* double mutant (Cavalier et al. 2008) and *xxt3xxt4xxt5* triple mutant (Zhang et al. 2023) were obtained earlier. To observe the root hair phenotype, sterilized seeds were grown on Petri dishes in ½ MS media (pH 5.8) with 1% (w/v) agar under long-day conditions (16 h: 8 h, light: dark) at 22°C. Seeds were germinated in horizontally positioned dishes, which were turned vertically later to ensure the roots grew straight.

To generate multiple mutant XXTs, the gene-specific primers (Supplemental Table S1) were utilized to clone the two fragments of truncated XXTs from previous vectors (Chou et al. 2012, 2015), and the fusion PCR was applied to link the two fragments as the full length of truncated XXTs. The Hot Start II DNA Polymerase (Thermo Fisher F122) was used in the fusion PCR: 5X Phire Reaction Buffer 10 μL; forward primer (2 µM) 12.5 μL; reverse primer (2 µM) 12.5 μL; 10 mM dNTPs 1 μL; DMSO 1.5 μL; Fragment 1(20 ng/ μL) 2 μL; Fragment 2 (20 ng/ μL) 2 μL; Phire Hot Start II DNA Polymerase 1 μL and add double-distilled water (ddH2O) to 50 μL. The Cycling steps were 98 ℃ for 5 minutes, followed by 35 cycles (98 ℃ for 10 seconds, 70 ℃ for 30 seconds, 67 ℃ for 30 seconds, 64 ℃ for 30 seconds, and 72 ℃ for 1 minute), then 72 ℃ for 5 minutes, and finally 12 ℃ for ∞. The fusion PCR products were digested with the corresponding restriction enzymes and inserted into pUBN-CFP expression vectors. For the BiFC assay and confocal microscopy study, all genes (AtXXT2, AtXXT5, AtSar1d, AtSec24a and α-Mannosidase (MNS1)) were cloned with gene-specific primers from previous vectors (Chou et al., 2012, 2015) and full-length complementary (cDNA). The PCR products were digested with the corresponding restriction enzymes and inserted into either the N- or C-terminal fragment of YFP (nYFP, cYFP) at the N-terminal (Chou et al. 2012, 2015) in pSAT vectors (Citovsky et al. 2006).

### Transgenic *Arabidopsis* plants

Recombinant *Agrobacterium* with pUBN::CFP-XXT2 (XXT2) or pUBN::CFP-XXT2-RQRQ (XXT2-RQRQ) was transformed into the *xxt1xxt2* double mutant, and *Agrobacterium* with pUBN::CFP-XXT5 (XXT5) or pUBN::CFP-XXT5-RQRQ (XXT5-RQRQ) was transformed into the *xxt3xxt4xxt5* triple mutant by the floral dipping method (Clough and Bent 1998). Sterilized transformants were selected on the Petri dishes with 250 µg/mL Hygromycin B (Thermo Fisher 10687010), and the expression of CFP in the transgenic seedlings was confirmed by detecting the CFP fluorescence under the fluorescent microscope. The protein expression levels of CFP–XXT2, CFP–XXT2-RQRQ, CFP–XXT5, and CFP–XXT5-RQRQ were examined by western blot analysis (Figure S12).

### Transient expression in *Arabidopsis* protoplasts

The recombinant pUBN::CFP-XXTs, pUBN::CFP-truncated XXTs (including XXT2Δ12A; XXT2Δ15A; XXT2ΔN; XXT2ΔNM; XXT2ΔM; XXT5Δ10A; XXT5ΔTTT; XXT5Δdi-Arg; XXT5Δ40A and XXT5ΔN) and pUBN::CFP-mutated XXTs (including XXT2-1RQ; XXT2-2RQ; XXT2-RQRQ; XXT5-RGGR; XXT5-RGR and XXT5-RQRQ) vectors were extracted from *E. coli* cells and were purified via PureLink™ HiPure Plasmid Maxiprep Kit (Invitrogen K210007). The preparation of *Arabidopsis* protoplasts from Col-0 and transgenic *Arabidopsis* and the t, as well as the transient expression of constructs, was performed according to a previously described (Chou et al. 2012, 2015; Zhang et al. 2023). After overnight incubation in the dark at room temperature, the fluorescence signals of CFP, YFP, and mCherry were detected using a confocal microscope (Leica SP5 system and Leica STED super-resolution) (Zhang et al. 2023).

For the BiFC assay, pSAT vectors with different genes were purified and transformed into *Arabidopsis* protoplasts. After 2 days of incubation in the dark at room temperature, fluorescence signals were checked with the same confocal (Chou et al. 2012, 2015).

### Cell wall extraction and analysis of hemicellulose fractions

Alcohol-insoluble cell wall material and the hemicellulose fractions from all studied plant lines and Col-0 were extracted using established standard protocols (Zabotina et al. 2012; Zhang et al. 2023). The hemicellulose fractions digested with endo-β-1,4-glucanase (XEG) were analyzed by matrix-assisted laser desorption ionization time-of-flight (MALDI-TOF) mass spectrometry. The protocol followed was published in (Zhang et al. 2023). The same hemicellulose fractions were separately digested with Driselase, a mixture of exoglycosidases and endoglycosidases (Sigma-Aldrich, D9515), and the total amount of a released signature disaccharide, isoprimeverose [IP; Xyl-a-(1-6)-Glc] was quantified by HPAEC using the gradient conditions described previously (Zabotina et al. 2012; Zhang et al. 2023).

### Protein extraction

About 15g of 14-day-old young *Arabidopsis* seedlings expressing CFP-XXT2 and CFP-XXT5 were collected and ground in liquid Nitrogen. A total of 60 mL protein extraction buffer (50 mM HEPES, 0.3 M sucrose, 65 mM NaCl, 3 mM EDTA, pH 7.5) supplemented with 3 mL protease inhibitor cocktail (1 mM E-64, 1 mM leupeptin, 100 mM AEBSF, 100 mM benzamidine, and 100 mM PMSF in methanol) was added. Plant debris was removed by filtration through Miracloth, and the clarified filtration was transferred to new 50 mL centrifuge tubes. The filtration was centrifuged at 20,000 × g for 1 h at 4°C, and the supernatant was transferred to ultracentrifuge tubes (Beckman #355618). The supernatant was then ultracentrifuged at 100,000 × g for 1 h at 4°C using a Ti70 rotor. Then the supernatant was discarded, and the pellet was washed with 1 mL protein extraction buffer. After the three washes, the membrane pellet was resuspended in 500 µL of protein extraction buffer containing 2% Triton X-100 and 1% NP-40, and incubated with shaking at 4°C overnight to solubilize the membrane-bound proteins. The solubilized membrane fraction was transferred to 1.5 mL tubes, kept on ice, and sonicated to further enhance solubilization. Finally, the samples were centrifuged at 14,000 × g 10 mins and 4°C to remove the insoluble remaining aggregation, and the supernatants (containing total membrane protein) were collected for subsequent analysis.

### Protein purification

The cDNA of the COPII complex proteins AtSar1b (AT1G56330), AtSar1c (AT4G02080), AtSar1d (AT3G62560) and AtSec24a (AT3G07100) were inserted into the pET His6 Sumo TEV LIC cloning vector containing an N-terminal 6xHis tag using gene-specific forward and reverse primers and expressed in the *E. coli* (BL21). Transformed *E. coli* was incubated in 500 mL of lysogeny broth medium at 37 °C at 180 rpm, and when cells reached a density with OD of 0.4-0.6, the temperature was reduced to 18 °C, isopropyl-β-D-thiogalactopyranoside (IPTG) was added to a final concentration of 0.5 mM, and the cells were incubated for another 18 hours at 18°C. Collected cells were resuspended in the cold lysis buffer (25 mM Tris; pH 7.4, 300 mM NaCl, 0.5 mM EDTA), and treated by five freeze/thaw cycles in liquid nitrogen. After thawing, the lysozyme solution was added to achieve a final concentration of 1 mg/mL, and the cells were incubated at 4 °C for 30 minutes to 60 minutes. The cells were sonicated five times for 15 seconds each and centrifuged at 20,000 × g for 30-60 minutes to collect soluble proteins. The soluble protein fractions were passed through HisPur Ni-NTA Resin (Thermo Scientific, 88223) to purify the recombinant proteins, following the manufacturer’s protocol. Amicon Ultra-15 Centrifugal Filter units (Millipore Sigma, UFC901024 and UFC903024) were used to purify the proteins further and to replace the HisPur™ Ni-NTA Resin elution buffer (50 mM Tris-HCl, 150 mM NaCl, 300 mM Imidazole) to the lysis buffer (25 mM Tris; pH 7.4, 300 mM NaCl, 0.5 mM EDTA). The purified COPII complex proteins AtSar1b, AtSar1c, AtSar1d, and AtSec24a were checked by the Coomassie stains and Western blot (Figure S13). Purified proteins were stored in the lysis buffer and were used for the pull-down assay *in vitro*.

### *In Vitro* Pull-Down Assay

In *vitro* pull-down assays with full-length XXT2 and XXT5, 400μg Sar1 or Sec24a purified proteins, incubate with 100 μL of HisPur™ Ni-NTA Resin (Thermo Scientific, 88223) in each reaction at 4 °C for 1 hour with end-to-end shaking. After incubation, the resin in the column was washed five times with Wash Buffer (pH 7.4, 20 mM sodium phosphate, 300 mM sodium chloride, and 25 mM imidazole). After washing, the resin was incubated with 500 µL total membrane protein solution at 4 °C for 4 hours with end-to-end shaking. The same washing steps were repeated as described above, and proteins were eluted with 160 μL acid elution buffer (pH 7.4, 20 mM sodium phosphate, 300 mM sodium chloride, and 250 mM imidazole). Collected elution fractions were analyzed by western blotting using tag-specific antibodies. Full length of XXT2 and XXT5 proteins were individually incubated with HisPur™ Ni-NTA Resin as the negative control, and the results are shown in Figure S14.

The cytosolic N-end tails of XXT2, XXT5 fused with Flag-tag were synthesized by PEPTIDE 2.0 Inc. The sequences of the synthesized peptides: **XXT2:** MIERCLGAYRCRRIQRALRQLKDYKDDDDK, **XXT2Q:** MIERCLGAYRCRRIQQALQQLKDYKDDDDK, **XXT2A:** MIERCLGAYRCRRIQAALAQLKDYKDDDDK.

**XXT5:** MGQDGSPAHKRPSGSGGGLPTTTLTNGGGRGGRGGLLPRGRQMQKTFNNIKDYKD DDDK,

**XXT5Q:** MGQDGSPAHKRPSGSGGGLPTTTLTNGGGQGGQGGLLPQGQQMQKTFNNIKDYKD DDDK,

**XXT5A:** MGQDGSPAHKRPSGSGGGLPTTTLTNGGGAGGAGGLLPAGAQMQKTFNNIKDYKDD DDK.

In *in vitro* pull-down assays with cytosolic N-end tails of XXT2 and XXT5, the Pierce Anti-DYKDDDDK Affinity Resin (Thermo Scientific, PIA36801), and the company protocol were used. For each reaction, 50 μL of resin was washed three times with cold lysis buffer (25 mM Tris, pH 7.4, 300 mM NaCl, 0.5 mM EDTA) and incubated with 40 μg of peptides with a flag-tag dissolved in 300 μL of lysis buffer at 4 °C for 4 hours with end-to-end shaking. After incubation, the resin in the column was washed twice with PBS (Phosphate Buffered Saline) buffer and once with ddH2O. After washing, the resin was incubated with 4 nmol of each COPII complex protein, dissolved in lysis buffer, at 4 °C overnight with end-to-end shaking. The same washing steps were repeated as described above, and proteins were eluted with 200 μL of acid elution buffer (0.1 M glycine; pH 2.8). The acid was then immediately neutralized by adding neutralization buffer (1 M Tris; pH 8.5). Collected elution fractions were analyzed by western blotting using tag-specific antibodies. AtSar1 and AtSec24a proteins were individually incubated with Pierce Anti-DYKDDDDK Affinity Resin as the negative control, and the results are shown in Figure S15.

### Western Blotting

The SDS-PAGE with an 18% acrylamide gel was used for the separation of the peptides with a Flag tag, while a 10% acrylamide gel was used for the separation of AtSar1, and 8% acrylamide gel was used for the separation of XXTs and AtSec24A recombinant proteins. After SDS-PAGE separation, the proteins were transferred to nitrocellulose membranes (0.2 mm; Bio-Rad) for immunodetection. DYKDDDDK Tag monoclonal antibodies (MA191878; Invitrogen) were used (1:1000 dilution) to detect synthesized XXTs peptides. Anti-Myc Tag antibodies (Sino Biological; 50-161-0309) were used (1:2000 dilution) to detect the AtSar1 proteins. HA Tag monoclonal antibodies (Invitrogen; PI26183) were used (1:2000 dilution) to detect the AtSec24A proteins. Monoclonal anti-GFP antibodies (MMS-118P; Covance) were used (1:6000 dilution) to detect the CFP-XXTs fusion proteins. The membranes were incubated with West Pico PLUS Chemiluminescent Substrate (SuperSignal) and visualized using a ChemiDocXRS+ (Bio-Rad) system. The intensities of the bands on the membrane were quantified using ImageJ.

## Supplemental Data

Supplemental Table. S1 The primers used in the study.

Supplemental Figure. S1 Support Figure 1. The subcellular localization of truncated XXT2 and XXT5 in the *Arabidopsis* protoplasts.

Supplemental Figure. S2 The bright field of the *Arabidopsis* protoplasts.

Supplemental Figure. S3 The bright field of the *Arabidopsis* protoplasts.

Supplemental Figure. S4 The subcellular localization of mutant XXT2 and XXT5 in the *Arabidopsis* protoplasts.

Supplemental Figure. S5 The bright field of the *Arabidopsis* protoplasts.

Supplemental Figure. S6 The bright field of the *Arabidopsis* protoplasts.

Supplemental Figure. S7 The subcellular localization of the wild type and mutated XXTs stably expressed in the *Arabidopsis* mutant plants.

Supplemental Figure. S8 The bright field of the *Arabidopsis* protoplasts prepared from stable expression lines.

Supplemental Figure. S9 The bright field of the *Arabidopsis* protoplasts prepared from stable expression lines.

Supplemental Figure. S10 Fluorescence images of BiFC signal in *Arabidopsis* protoplasts.

Supplemental Figure. S11 The protein-protein interactions between cytosolic tails of XXTs with AtSec24A proteins via pull-down assay in vitro.

Supplemental Figure. S12 The protein-protein interactions between cytosolic tails of XXTs with AtSec24A proteins via pull-down assay in vitro.

Supplemental Figure. S13 Coomassie Blue image and the corresponded western blot result of the protein purification of AtSar1 and AtSec24A proteins.

Supplemental Figure. S14 AtSar1 and AtSec24A proteins showed no binding to the anti-Flag resin.

Supplemental Figure. S15 AtXXT2 and AtXXT5 proteins showed no binds to the anti-His resin.

## Acknowledgments

This study was supported by NSF-MCB (grant #1856477). Open access funding is provided by the Iowa State University Library.

## Author contributions

OAZ conceived the research ideas and developed the project. NZ and JDJ did the experiments and collected the data. NZ wrote the manuscript. OAZ and NZ contributed to manuscript preparation.

## Data availability

All the data and materials that support the findings of this study are available upon request from the corresponding author.

