## supplemental table 1 for "The interaction of the Arabidopsis Xyloglucan Xylosyltransferases XXTs with the COPII member SAR1 via their di-Arginine motifs is critical for delivery to the Golgi"

**Table. S1 The primers used in the study.**

| Name | Sequence |
|---|---|
| CFP-XXT2-F | *ATACGCGTCGACATGATTGAGAGGTGTTTAG* |
| CFP-XXT2-R | *ATTACCCTGTTATCCCTATCAAACTTGATTGGTTTGTAC* |
| CFP-XXT2-12AA-F | *ATACGCGTCGACATGAGAATCCAAAGAGCTTTACG* |
| CFP-XXT2-15AA-F | *ATACGCGTCGACAGAGCTTTACGCCAATTAAAGGTAAC* |
| CFP-XXT2-N-F | *ATACGCGTCGACGTAACGATTCTCTGTC* |
| CFP-XXT2-NM-F | *ATACGCGTCGACGTTGTCCTACGTAGCAC* |
| CFP- XXT2-M-F | *ATACGCGTCGACAAATTCGGAACTCCGGAGC* |
| CFP-XXT2-R-65 | *ATTACCCTGTTATCCCTATCAAACTTGATTGGTTTGTACCTTAACCG* |
| XXT2-R-Q-1-F | *GGAGAATCCAACAAGCTTTACGCCAATTAAAG* |
| XXT2-R-Q-1-R | *ACCTTTAATTGGCGTAAAGCTTGTTGGATTCTC* |
| XXT2-R-Q-2-F | *CAAAGAGCTTTACAGCAATTAAAGGTAACGATTCTC* |
| XXT2-R-Q-2-R | *CGTTACCTTTAATTGCTGTAAAGCTCTTTGGA* |
| XXT2-RQRQ-1,2-F | *GAGAATCCAACAAGCTTTACAGCAATTAAAGGT* |
| XXT2-RQRQ-1,2-R | *ACCTTTAATTGCTGTAAAGCTTGTTGGATTCTC* |
| CFP-XXT5-F | *ATACGCGTCGACATGGGTCAAGATGGTTCG* |
| CFP-XXT5-R | *ATTACCCTGTTATCCCTACTAGTTCTGTGGTTTGGTTTCCAC* |
| CFP-XXT5-10AA-F | *ATACGCGTCGACAGACCATCCGGAAGCGGC* |
| CFP-XXT5-TTT-F | *ATACGCGTCGACCTTCCGACGACGAC* |
| CFP- XXT5-di-Arg-F | *ATACGCGTCGACGGTGGAGGAAGAGGTGGT* |
| CFP-XXT5-40AA-F | *ATACGCGTCGACAGACAGATGCAGAAGACGTTTAAC* |
| CFP-XXT5-N-F | *ATACGCGTCGACATCACAATTCTCTGTGGTTTCGT* |
| XXT5-RQ-RGGR-F | *GGTGGAGGACAAGGTGGTCAGGGTGGT* |
| XXT5-RQ-RGGR-R | *ACCACCCTGACCACCTTGTCCTCCACC* |
| XXT5-RQ-RGR-F | *ACCTCAAGGTCAACAGATGCAGAAGACGTTTAACAACA* |
| XXT5-RQ-RGR-R | *TGTTGTTAAACGTCTTCTGCATCTGTTGACCTTGAGGT* |

| | |
|---|---|
| XXT5-RQRQ-F | *GAGGACAAGGTGGTCAGGGTGGTTTGTTACCTCAAGGTCAACAGATG* |
| XXT5-RQRQ-R | *CATCTGTTGACCTTGAGGTAACAAACCACCCTGACCACCTTGTCCTC* |
| 1092-XXT2-F | *CTTCGAATTCaATGATTGAGAGGTGTTTAGGAG* |
| 1092-XXT2-R | *GGTGGATCCTCAAACTTGATTGGTTTGTAC* |
| 1087-XXT5-F | *CTCGAGCTCAAGCTTCGAATATGGGTCAAGATGGTTCG* |
| 1087-XXT5-R | *GGTaGATCtTCAagcgtaatctggaacgtcatatggataGTTCTGTGGTTTGGTTTCC* |
| 1087-Sar1d-F | *GATCTCGAGCTCAAGCTTCGATGTTCCTGGTGGATTGG* |
| 1087-Sar1d-R | *GGTGGATCCagcgtaatctggaacgtcatatggataGTCGATATATTGAGAAACCCAT* |
| 1092-Sar1d-F | *CTTCGAATTCaATGTTCCTGGTGGATTGGT* |
| 1092-Sar1d-R | *GGTGGATCCTCAGTCGATATATTGAGAAACCCAT* |
| 1087-Sec24A-F | *GATCTCGAGCTCAAGCTTCGATGGGTACGGAGAATCAGGGC* |
| 1087- Sec24A-R | *GGTGGATCCagcgtaatctggaacgtcatatggataGTTTTGTTGAACTTGGCGGTGAA* |
| 1092- Sec24A-F | *GATCTCGAGatGTAATGGGTACGGAGAATCAGGG* |
| 1092- Sec24A-R | *AGGTGGATCCTCAGTTTTGTTGAACTTGGCGGTG* |
