## supplemental figure for "The interaction of the Arabidopsis Xyloglucan Xylosyltransferases XXTs with the COPII member SAR1 via their di-Arginine motifs is critical for delivery to the Golgi"

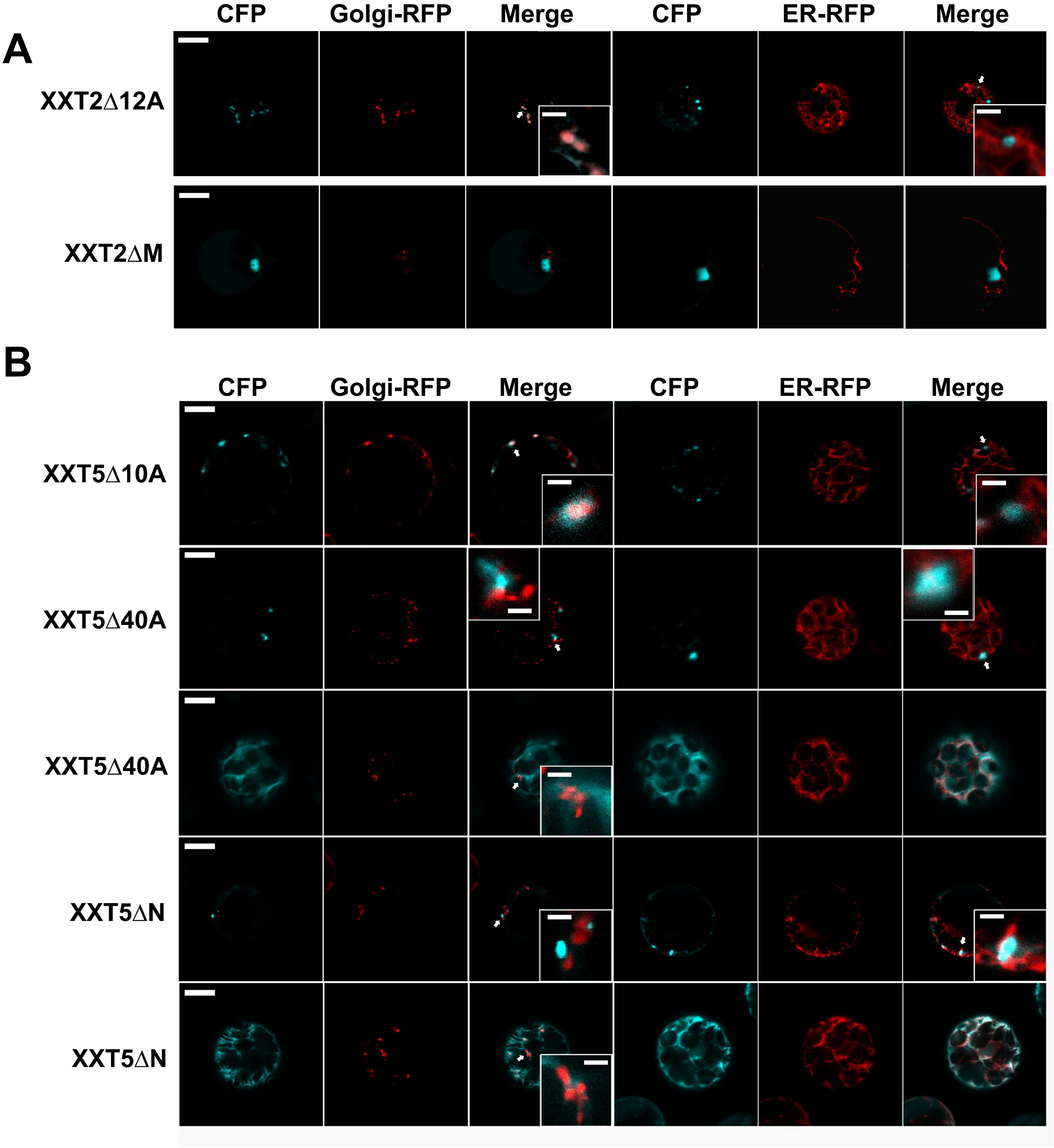
A
CFP
Golgi-RFP
Merge
CFP
ER-RFP
Merge
XXT2Δ12A
XXT2ΔM
B
CFP
Golgi-RFP
Merge
CFP
ER-RFP
Merge
XXT5Δ10A
XXT5Δ40A
XXT5Δ40A
XXT5ΔN
XXT5ΔN

**Figure S1. The subcellular localization of truncated XXT2 and XXT5 in the Arabidopsis protoplasts. A:** The subcellular localization of XXT2 and truncated mutants, co-expressed with Golgi-R (mCherry) or ER-R (mCherry) in Col-0 Arabidopsis protoplasts. The white arrows indicate Golgi localization of XXT2Δ12A. Bar, 10 μm. insets are shown at 4× magnification (scale bar = 2 μm). **B:** The subcellular localization of XXT5 and mutants, co-expressed with Golgi-R (mCherry) or ER-R (mCherry) in Col-0 Arabidopsis protoplasts. The white arrows indicate Golgi localization of XXT5Δ10A and non-Golgi-dot localization of XXT5Δ40A and XXT5ΔN. Bar, 10 μm. Insets are shown at 4× magnification (scale bar = 2 μm).

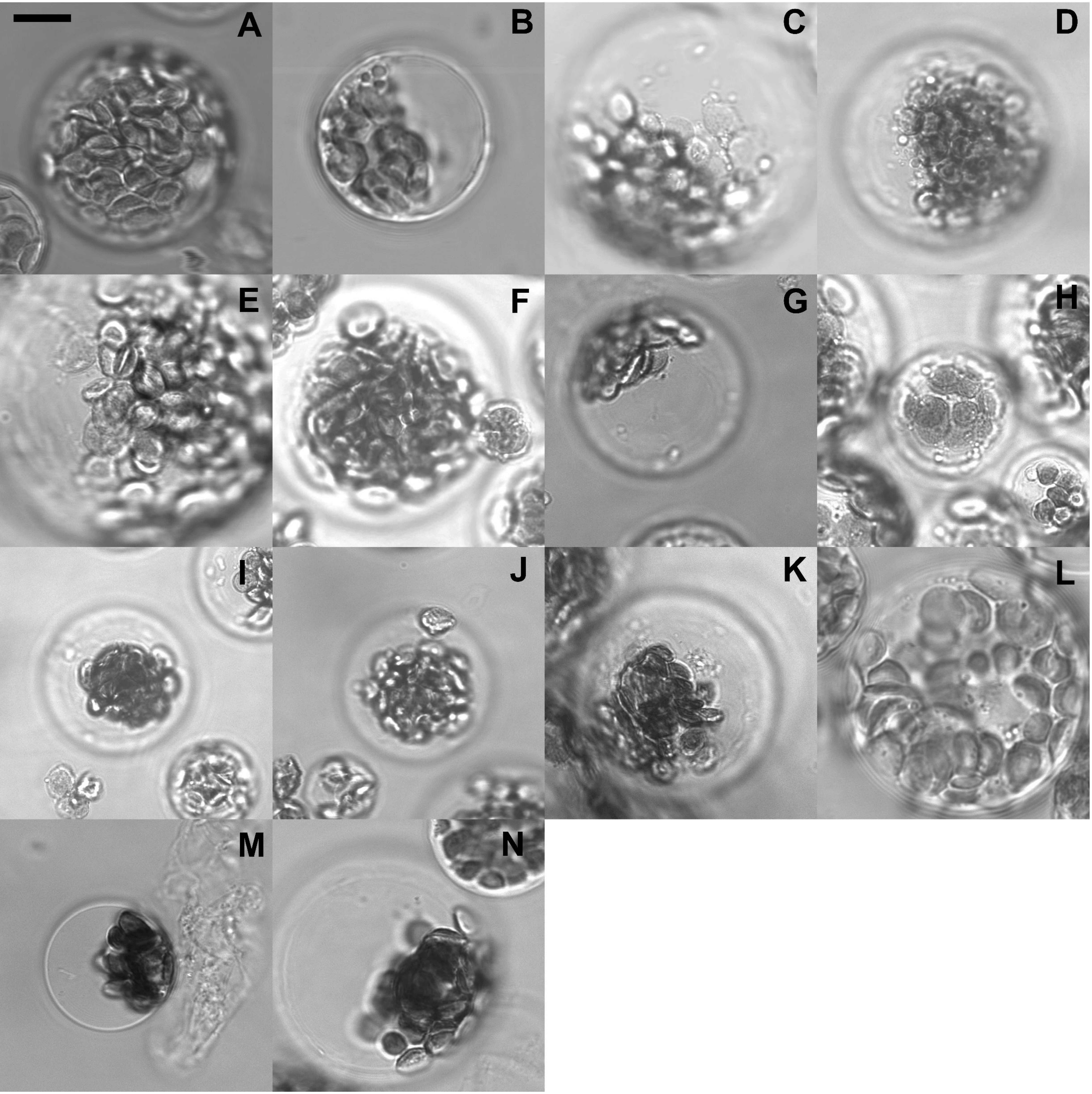
A
B
C
D
E
F
G
H
I
J
K
L
M
N

**Figure S2. The bright field of the Arabidopsis protoplasts. A** : XXT2 co-expressed with Golgi-R. **B**: XXT2 co-expressed with ER-R. **C** : XXT2Δ12A co-expressed with Golgi-R. **D**: XXT2Δ12A co-expressed with ER-R. **E**: XXT2Δ15A co-expressed with Golgi-R. **F**: XXT2Δ15A co-expressed with ER-R. **G, I**: XXT2ΔN co-expressed with Golgi-R. **H, J**: XXT2ΔN co-expressed with ER-R. **K**: XXT2ΔNM co-expressed with Golgi-R. **L**: XXT2ΔNM co-expressed with ER-R. **M**: XXT2ΔM co-expressed with Golgi-R. **N**: XXT2ΔM co-expressed with ER-R.

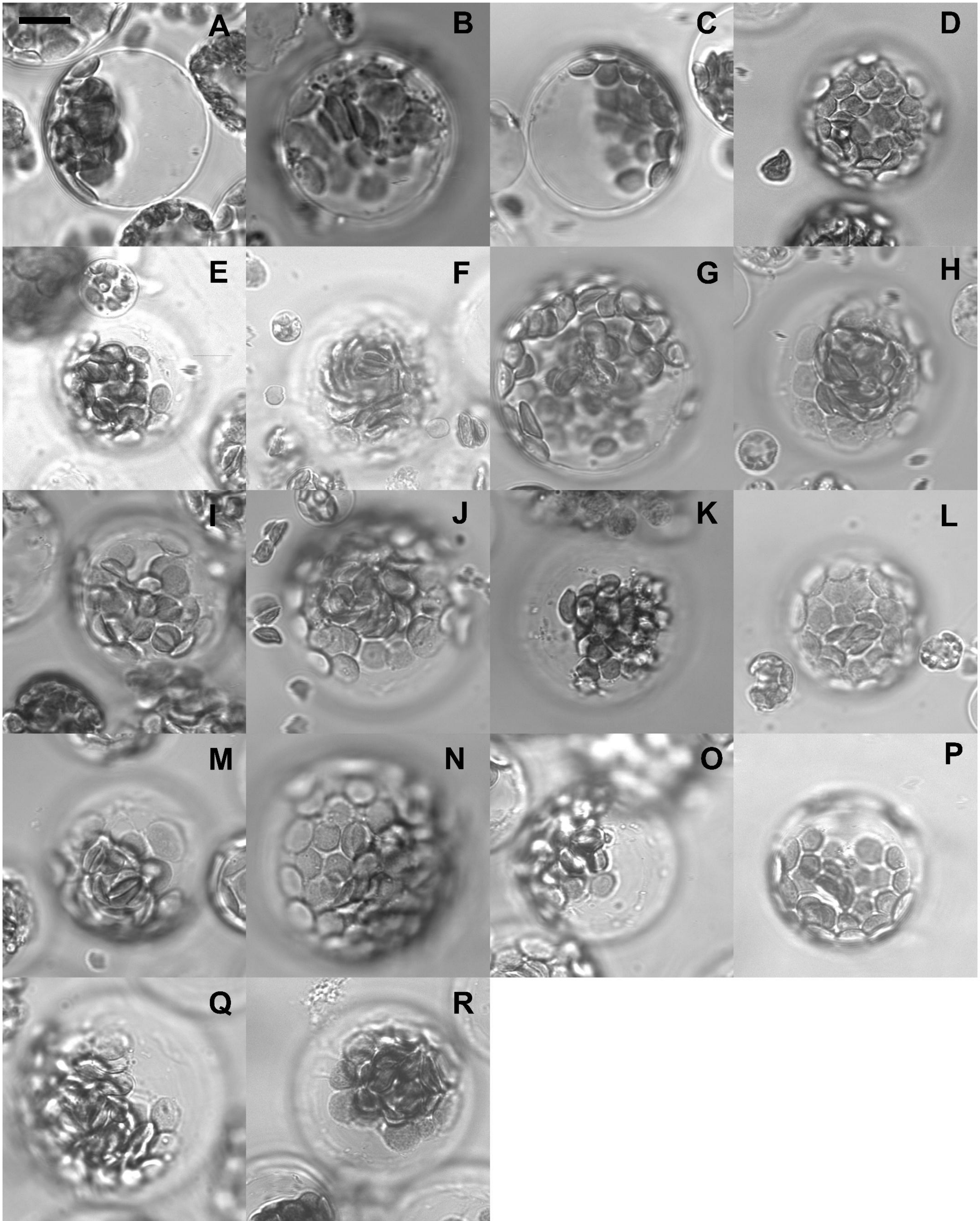

A
B
C
D
E
F
G
H
I
J
K
L
M
N
O
P
Q
R

**Figure S3. The bright field of the Arabidopsis protoplasts. A** : XXT5 co-expressed with Golgi-R. **B**: XXT5 co-expressed with ER-R. **C**: XXT5Δ10A co-expressed with Golgi-R. **D**: XXT5Δ10A co-expressed with ER-R. **E**: XXT5ΔTTT co-expressed with Golgi-R. **F**: XXT5ΔTTT co-expressed with ER-R. **G, I**: XXT5Δdi-Arg co-expressed with Golgi-R. **H, J**: XXT5Δdi-Arg co-expressed with ER-R. **K, M**: XXT5Δ40A co-expressed with Golgi-R. **L, N**: XXT5Δ40A co-expressed with ER-R. **O, Q**: XXT5ΔN co-expressed with Golgi-R. **P, R**: XXT5ΔN co-expressed with ER-R.

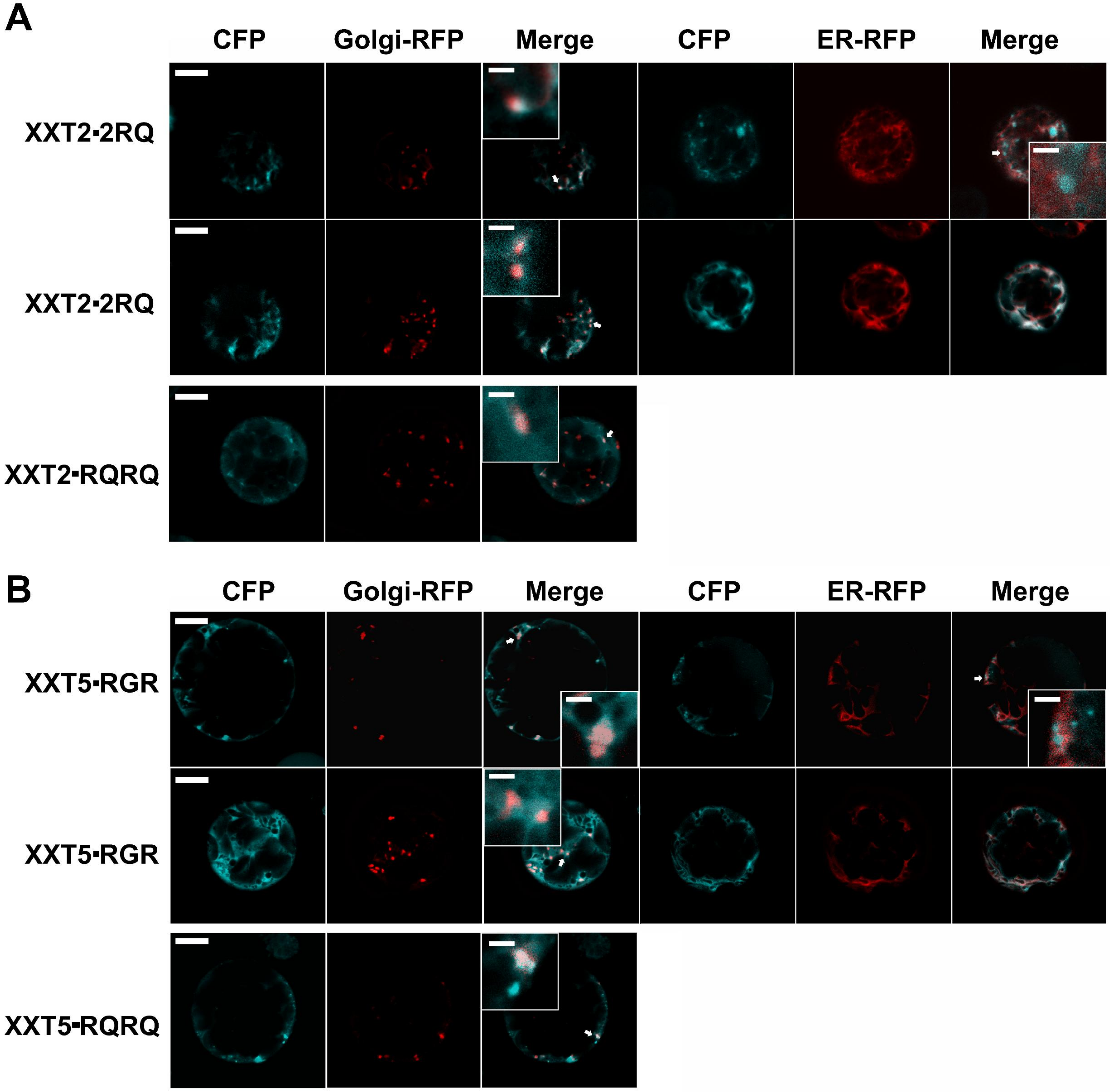
A
CFP
Golgi-RFP
Merge
CFP
ER-RFP
Merge
XXT2-2RQ
XXT2-2RQ
XXT2-RQRQ
B
CFP
Golgi-RFP
Merge
CFP
ER-RFP
Merge
XXT5-RGR
XXT5-RGR
XXT5-RQRQ

**Figure S4. The subcellular localization of mutant XXT2 and XXT5 in the Arabidopsis protoplasts. A:** The subcellular localization of mutant XXT2, co-expressed with Golgi-R (mCherry) or ER- R (mCherry) in Col-0 Arabidopsis protoplasts. The white arrows indicate Golgi localization of XXT2-2RQ and XXT2-RQRQ. Bar, 10 μm. Insets are shown at 4× magnification (scale bar = 2 μm). **B:** The subcellular localization of mutant XXT5, co-expressed with Golgi-R (mCherry) or ER-R (mCherry) in Col-0 Arabidopsis protoplasts. The white arrows indicate Golgi localization of XXT5-RGR and of XXT5-RQRQ. Bar, 10 μm. Insets are shown at 4× magnification (scale bar = 2 μm).

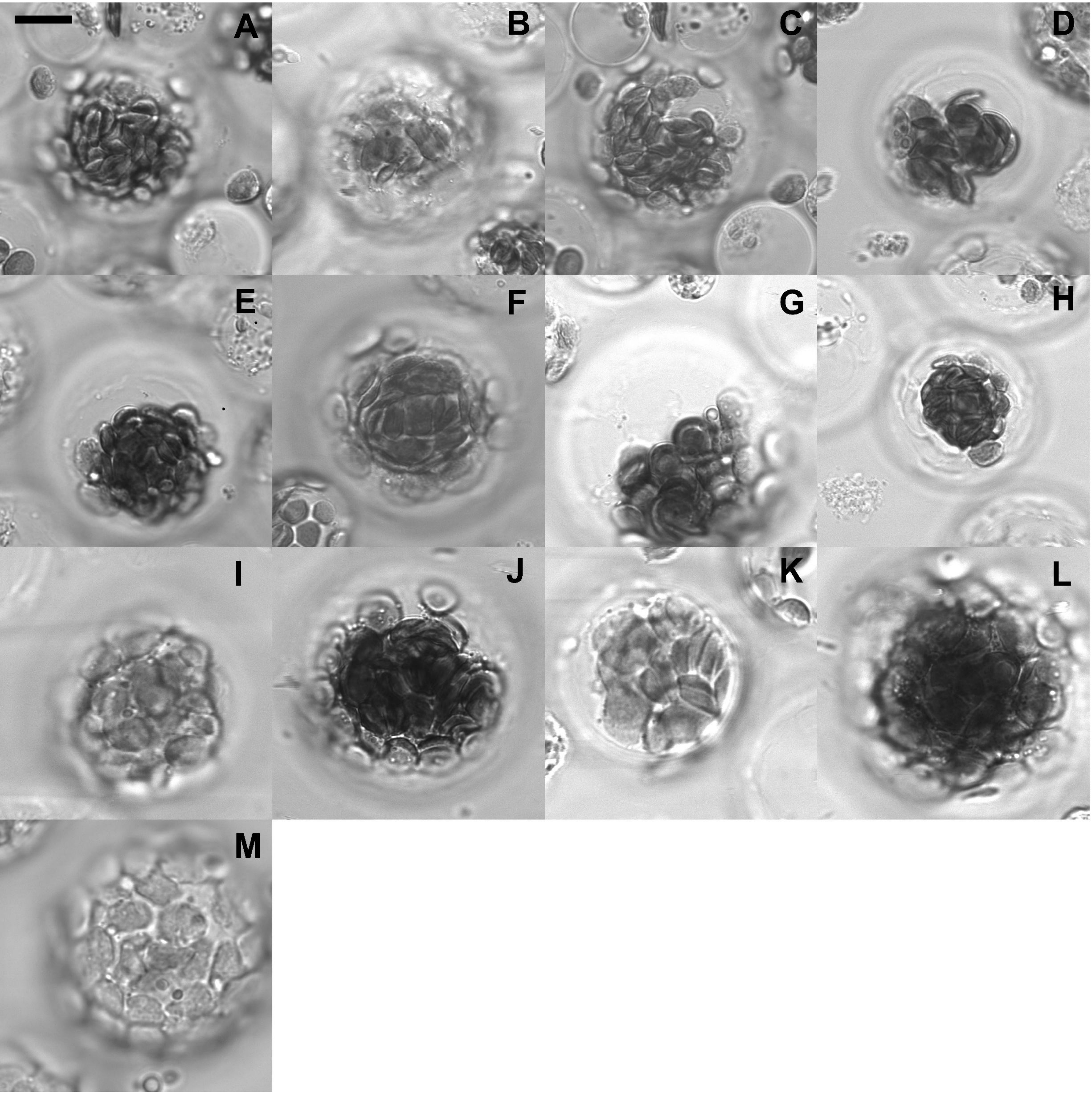
A
B
C
D
E
F
G
H
I
J
K
L
M

**Figure S5. The bright field of the *Arabidopsis* protoplasts. A, C**: XXT2-1RQ co-expressed with Golgi-R. **B, D**: XXT2-1RQ co-expressed with ER-R. **E, G**: XXT2-2RQ co-expressed with Golgi-R. **F, H**: XXT2-2RQ co-expressed with ER-R. **I, K, M**: XXT2-RQRQ co-expressed with Golgi-R. **J, L**: XXT2-RQRQ co-expressed with ER-R.

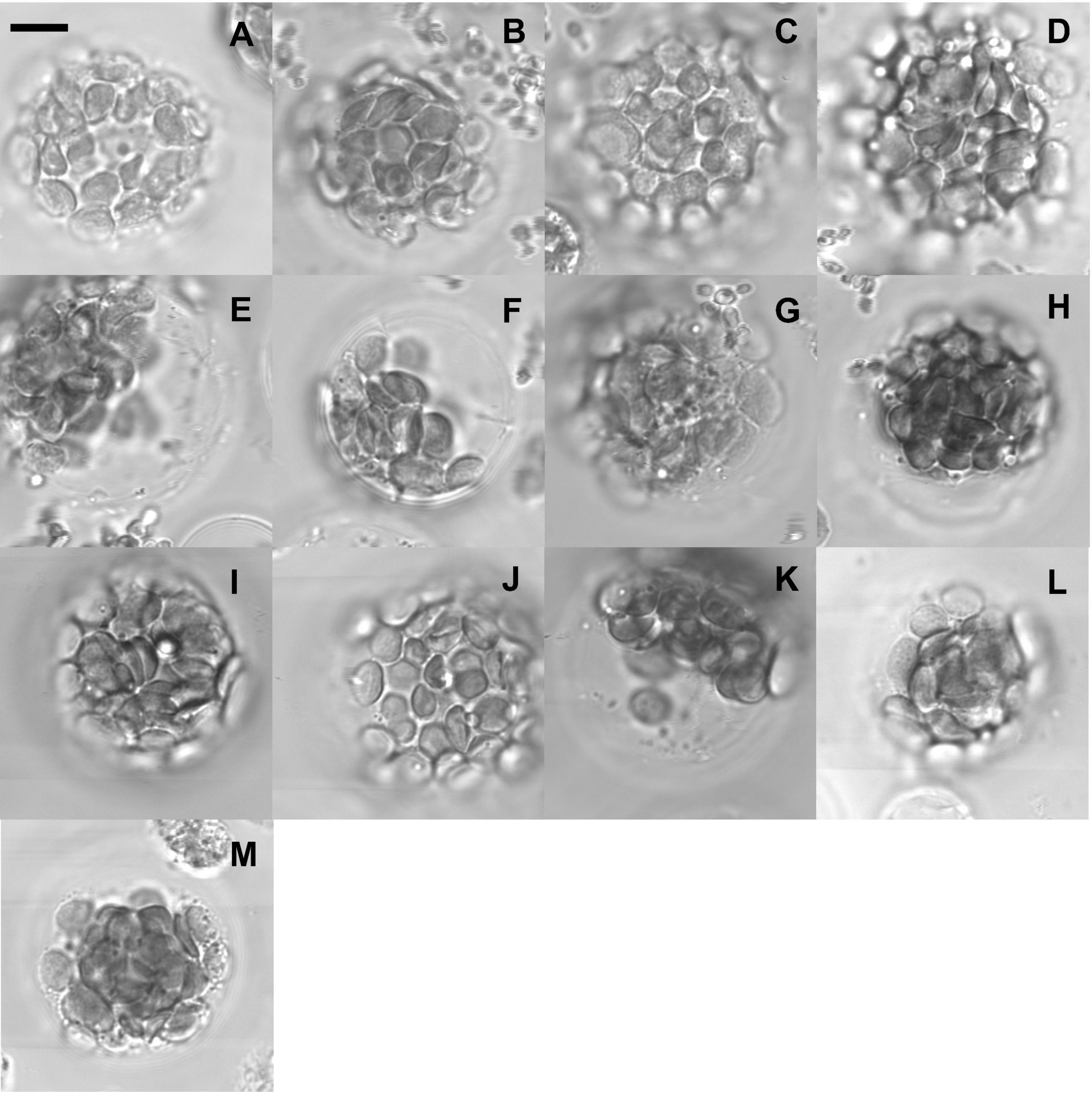

A
B
C
D
E
F
G
H
I
J
K
L
M

**Figure S6. The bright field of the *Arabidopsis* protoplasts. A, C**: XXT5 - RGGR co-expressed with Golgi-R. **B, D**: XXT5-RGGR co-expressed with ER-R. **E, G**: XXT5-RGR co-expressed with Golgi-R. **F, H**: XXT5-RGR co-expressed with ER-R. **I, K, M**: XXT5-RQRQ co-expressed with Golgi-R. **J, L**: XXT5-RQRQ co-expressed with ER-R.

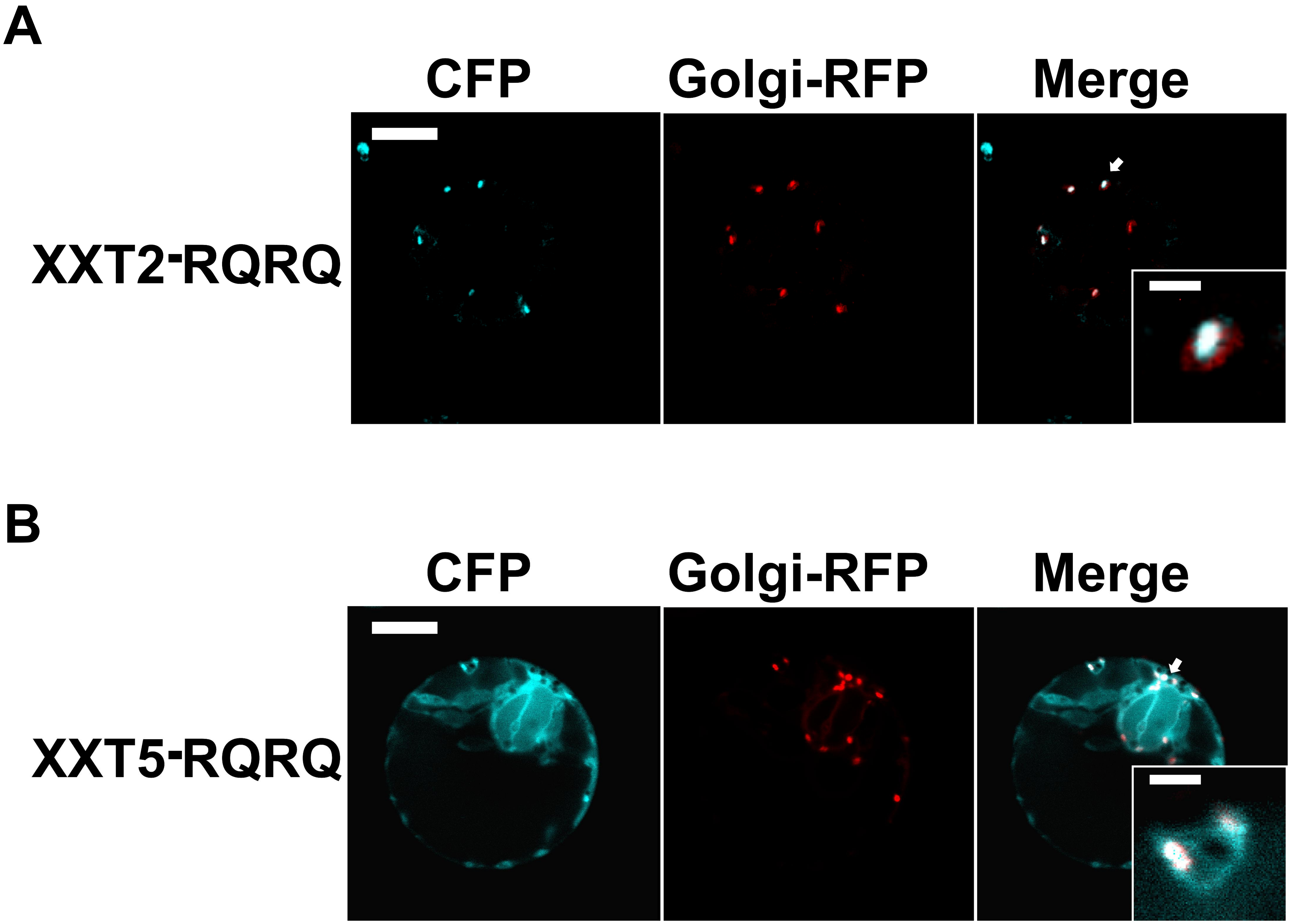
A
CFP
Golgi-RFP
Merge
XXT2-RQRQ
B
CFP
Golgi-RFP
Merge
XXT5-RQRQ

**Figure S7. The subcellular localization of the wild type and mutated XXTs stably expressed in the Arabidopsis mutant plants. A**: The subcellular localization of XXT2 RQRQ in the protoplasts prepared from the Arabidopsis *xxt1xxt2* plants expressing XXT2-RQRQ and transfected with Golgi R (mCherry)-R. Bar, 10 μm. Insets are shown at 4× magnification (scale bar = 2 μm). The white arrows indicate the Golgi localization of XXT2 -RQRQ. **B**: The subcellular localization of XXT5-RQRQ in the protoplasts prepared from the Arabidopsis *xxt3xxt4xxt5* plants expressing XXT5-RQRQ and transfected with Golgi-R (mCherry)-R. Bar, 10 μm. Insets are shown at 4× magnification (scale bar = 2 μm). The white arrows indicate the Golgi localization of XXT5-RQRQ.

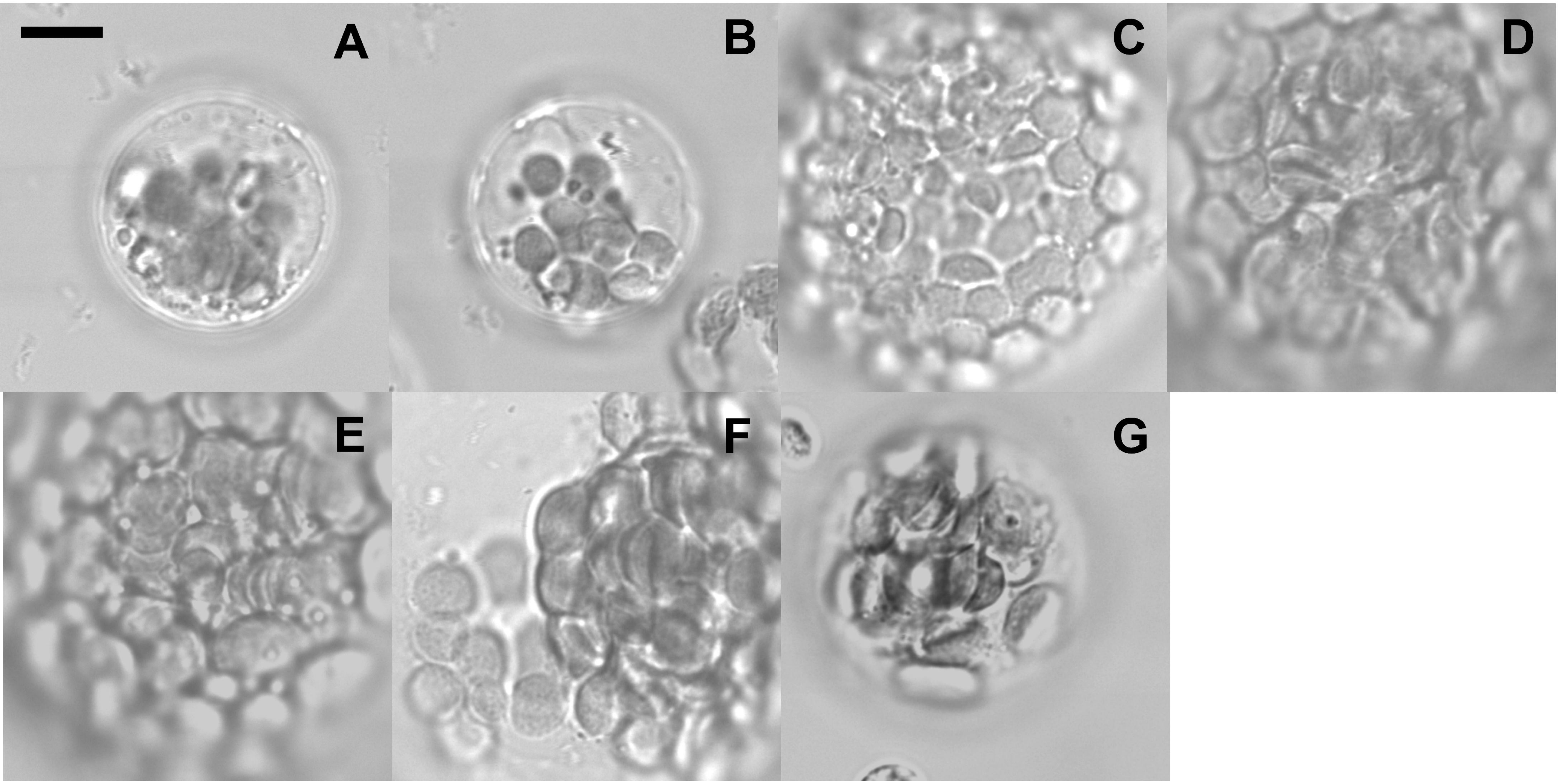


**Figure *S*8. The bright field of the *A*rabidopsis protoplasts prepared from stable expression lines. A**: expression of Golgi-R in the protoplast prepared from the stable expression of XXT2. **B**: expression of ER-R in the protoplast prepared from the stable expression of XXT2. **C, E, G:** : expression of Golgi-R in the protoplast prepared from the stable expression of XXT2-RQRQ. **D, F**: expression of ER-R in the protoplast prepared from the stable expression of XXT2-RQRQ

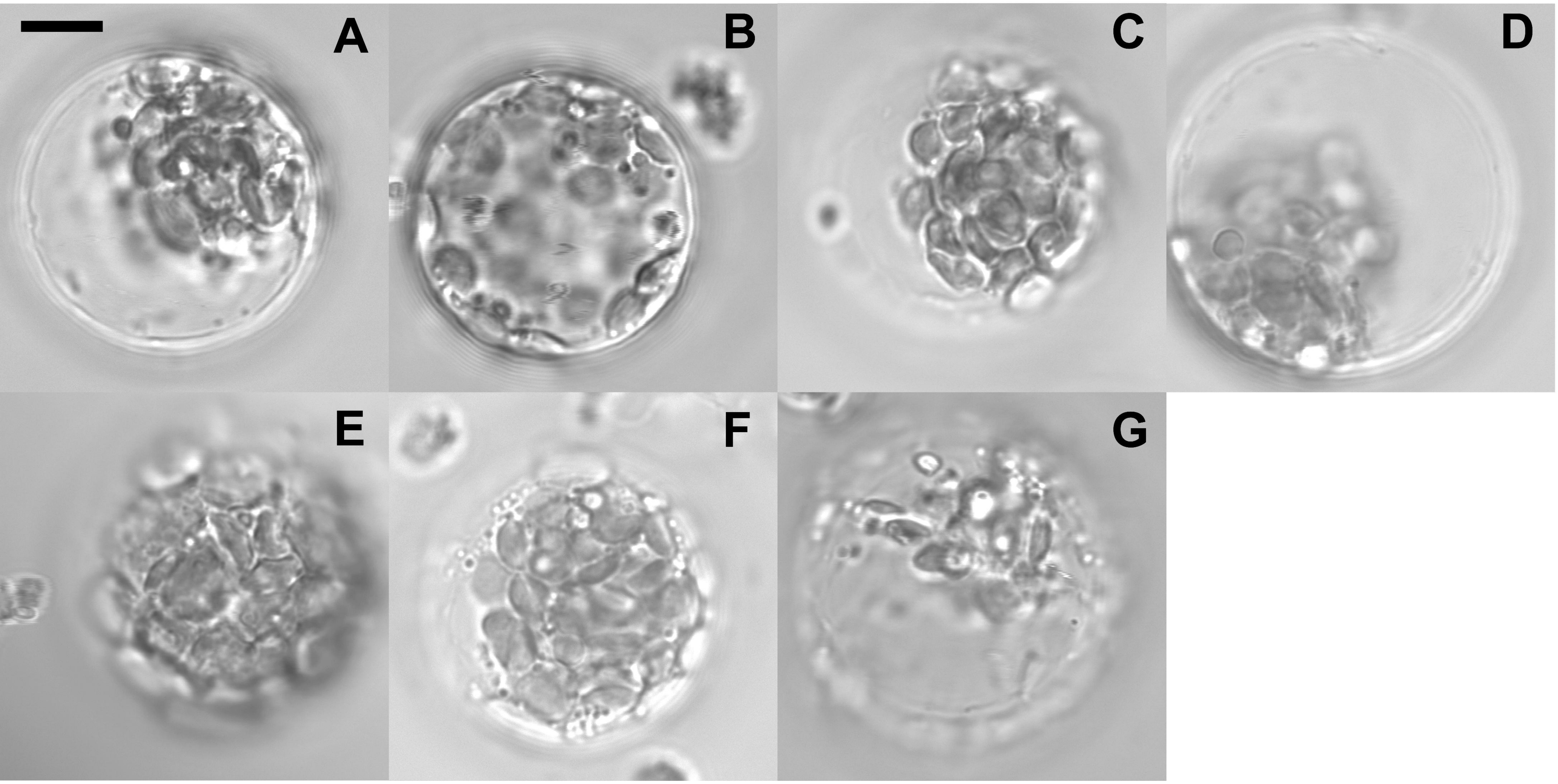


**Figure S9. The bright field of the *Arabidopsis* protoplasts prepared from stable expression lines. A**: expression of Golgi-R in the protoplast prepared from the stable expression of XXT5. **B**: expression of ER-R in the protoplast prepared from the stable expression of XXT5. **C, E, G:** : expression of Golgi-R in the protoplast prepared from the stable expression of XXT5-RQRQ. **D, F**: expression of ER-R in the protoplast prepared from the stable expression of XXT5-RQRQ

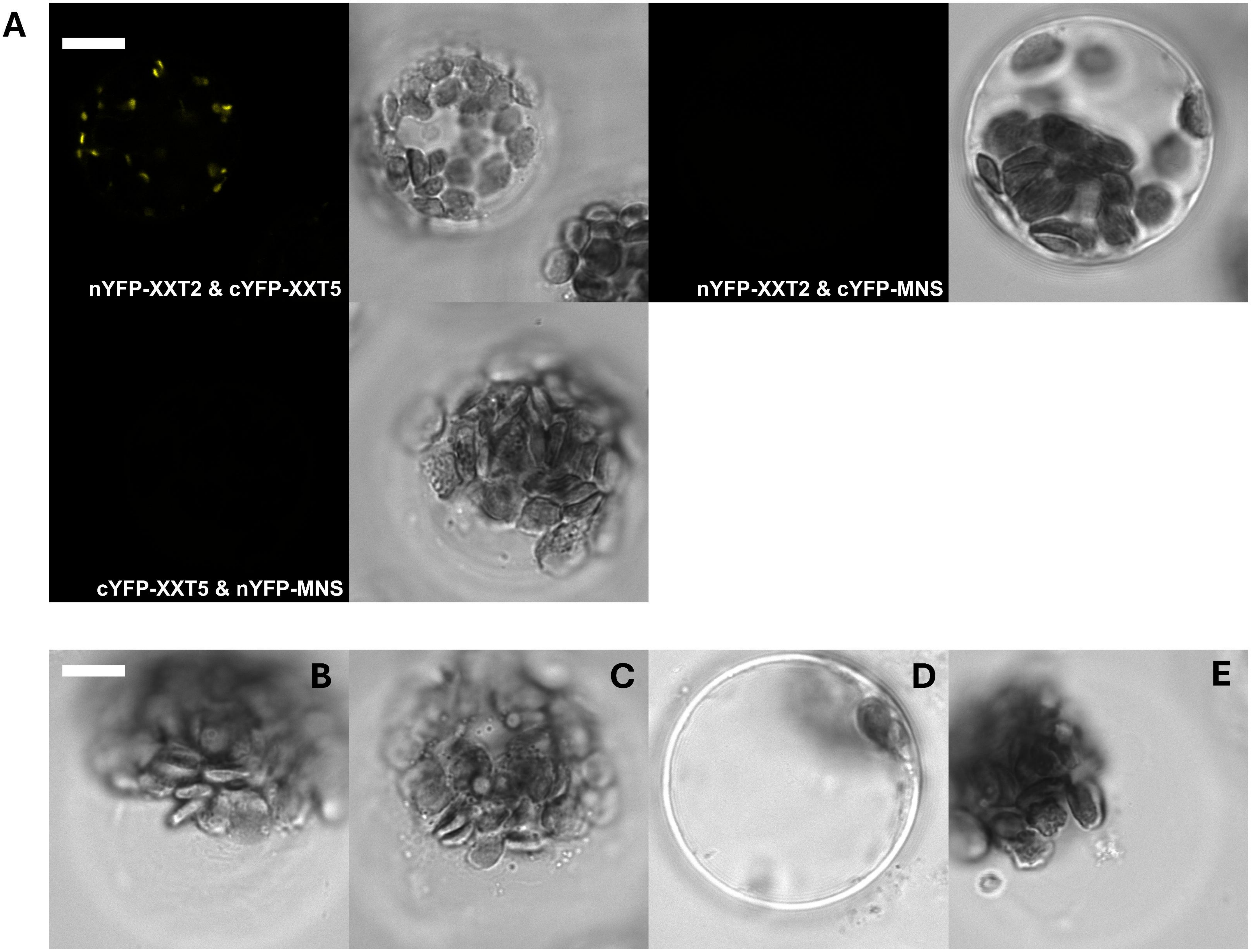
A
nYFP-XXT2 & cYFP-XXT5
nYFP-XXT2 & cYFP-MNS
cYFP-XXT5 & nYFP-MNS
B
C
D
E

**Figure S10. Fluorescence images of BiFC signal in Arabidopsis protoplasts. A:** fluorescence signal image and corresponding bright filed image. **B**: bright field image of BiFC pair nYFP-XXT2 & cYFP-Sar1d. **C**: bright field image of BiFC pair nYFP-XXT2 & cYFP-Sec24a. **D**: bright field image of BiFC pair cYFP-XXT5 & nYFP-Sar1d. **E**: bright field image of BiFC pair cYFP-XXT5 & nYFP-Sec24a. Bar, 10 μm.

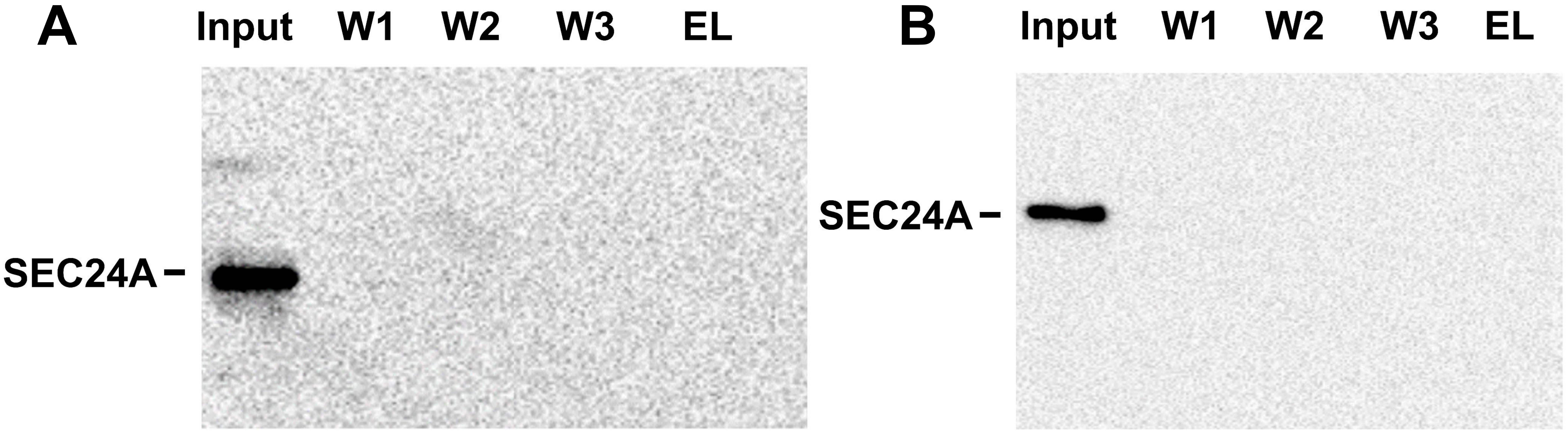


**Figure S11. The protein-protein interactions between cytosolic tails of XXTs with AtSec24a proteins via pull-down assay *in vitro*. A**: Detection of the protein-protein interactions between XXT2 with AtSec24a. **B**: Detection of the protein-protein interactions between XXT5 with AtSec24a.

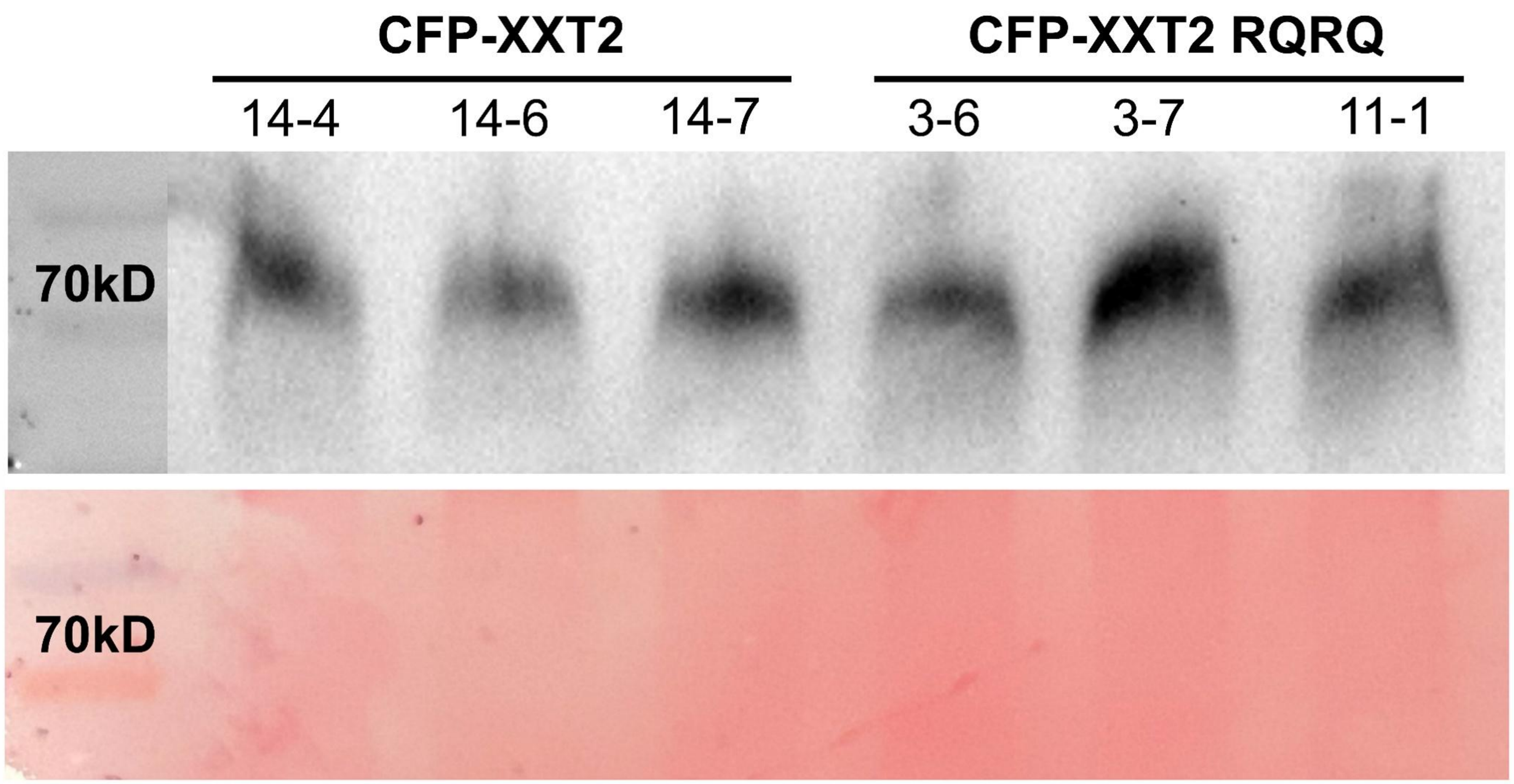
CFP-XXT2
CFP-XXT2 RQRQ
14-4
14-6
14-7
3-6
3-7
11-1
70kD
70kD

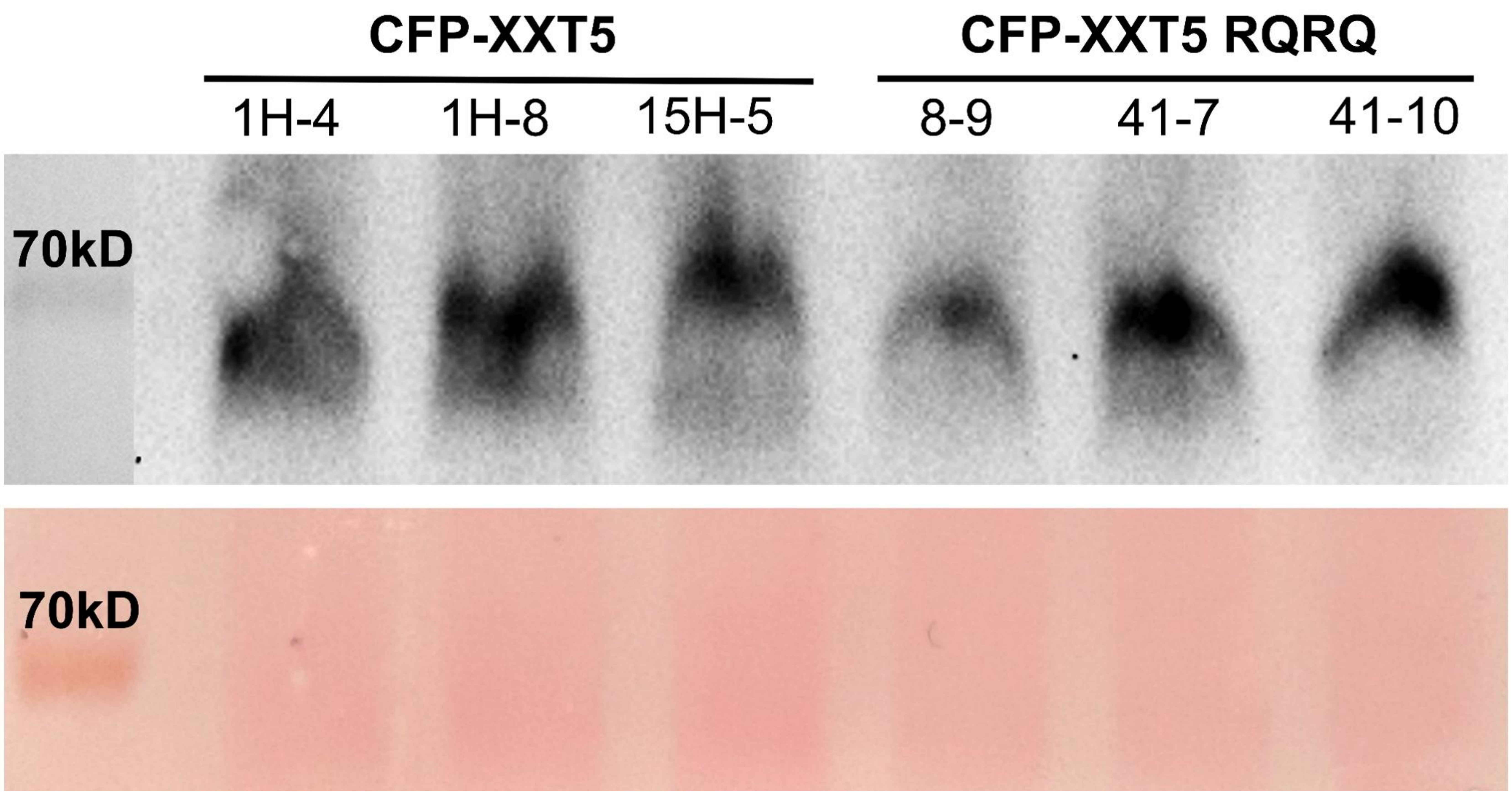
CFP-XXT5
CFP-XXT5 RQRQ
1H-4
1H-8
15H-5
8-9
41-7
41-10
70kD
70kD

**Figure S12. The protein expression level in the stable expression lines .** Detect the protein expression of CFP-XXT2, CFP-XXT2-RQRQ, CFP-XXT5 and CFP-XXT5-RQRQ in stable expression lines by western blot. Three independent lines are used in each group. Ponceau S is used to detect the all membrane protein.

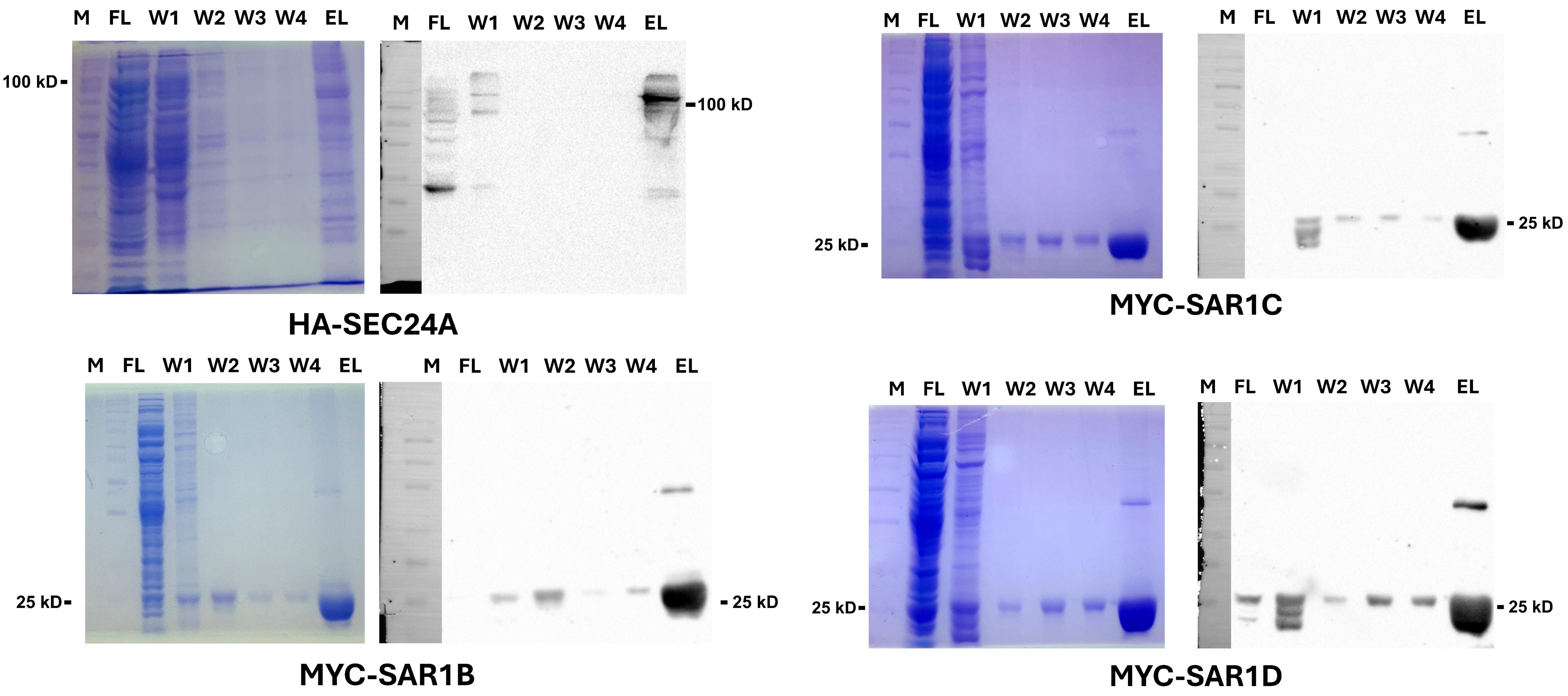


**Figure S13. Coomassie Blue image and corresponded western blot result of the protein purification of AtSar1 and AtSec24a proteins.** FT: flow through, W1: wash-1, W2: wash-2, W3: wash-3, W4: wash-4, W5: wash-5, EL: elution buffer.

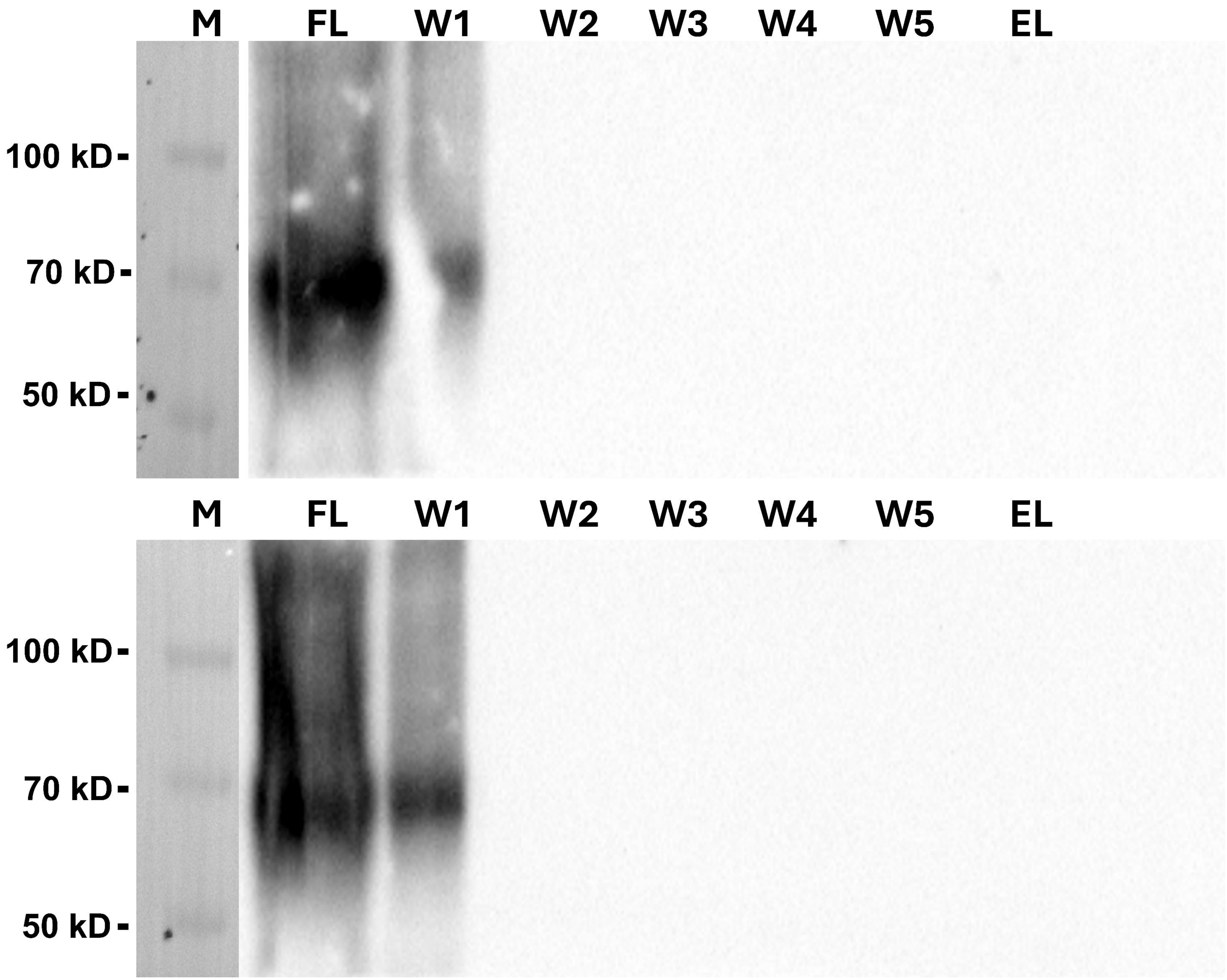


**Figure S14 AtXXT2 and AtXXT5 proteins showed no binds to the anti-His resin.** AtXXT2 and AtXXT5 had no binds to the anti-His resin. FT: flow through, W1: wash-1, W2: wash-2, W3: wash-3, W4: wash-4, W5: wash-5, EL: elution buffer.

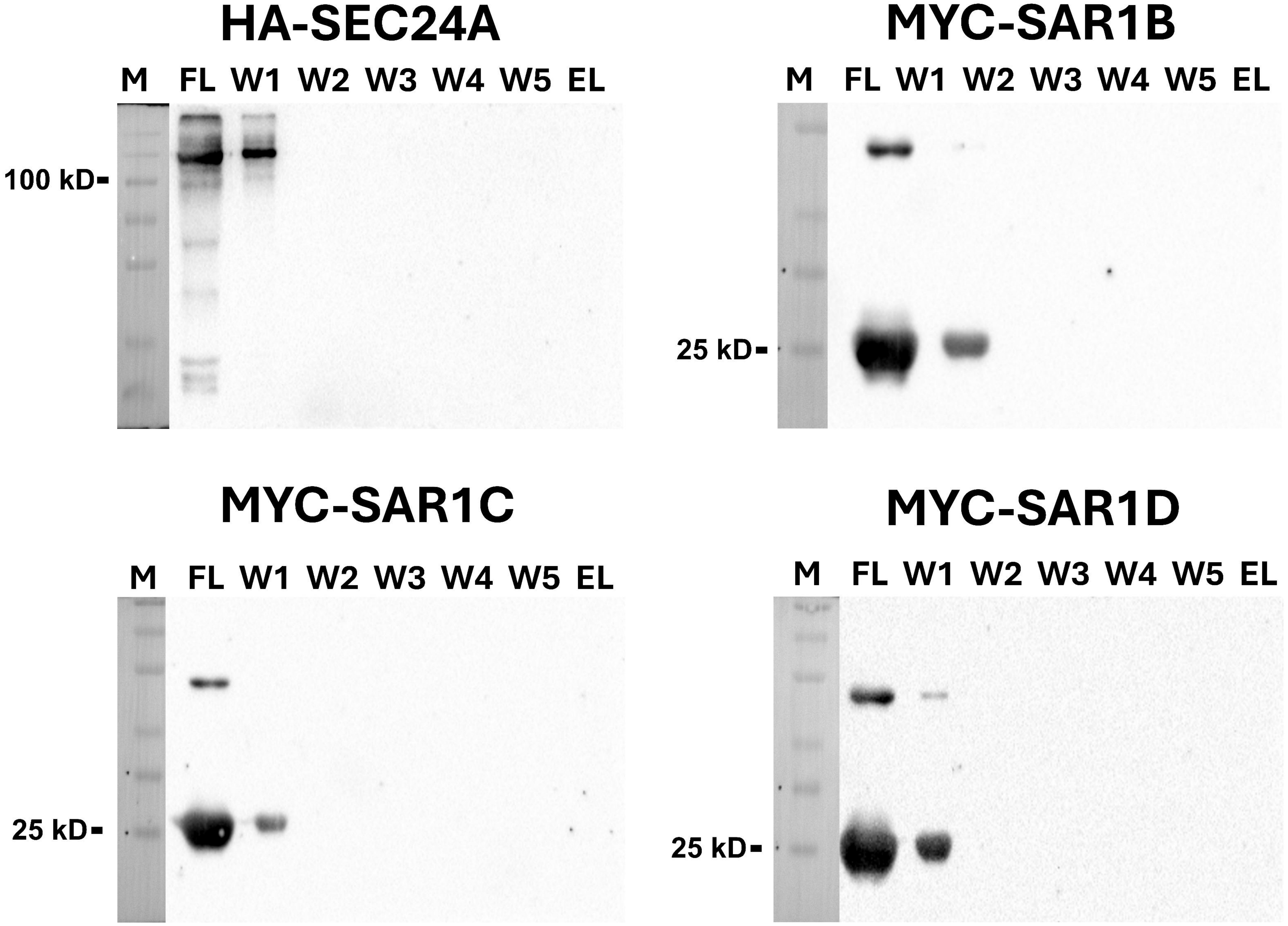


**Figure S15 AtSar1 and AtSec24a proteins showed no binds to the anti-Flag resin.** AtSar1b, AtSar1c, AtSar1d and AtSec24a had no binds to the anti-Flag resin. FT: flow through, W1: wash-1, W2: wash-2, W3: wash-3, W4: wash-4, W5: wash-5, EL: elution buffer.
